# Mutation-induced phenotypic plasticity drives formation of a pro-metastatic tumour-intrinsic niche

**DOI:** 10.1101/2022.12.22.521159

**Authors:** Jeroen M. Bugter, Layla El Bouazzaoui, Emre Küçükköse, Megan L. Mills, Yourae Hong, Joep Sprangers, Ingrid Jordens, Danai Kokkinidi, Susanna Plugge, Anton Venhuizen, Sabine Bosman, Joris Hageman, Nicola Fenderico, Kathryn Gilroy, Diego Montiel Gonzalez, Ruben van Boxtel, Saskia J.E. Suijkerbuijk, Sabine Tejpar, Andrew D. Campbell, Owen J. Sansom, Hugo J.G. Snippert, Onno Kranenburg, Madelon M. Maurice

## Abstract

Colorectal cancer (CRC) progression is characterised by a remarkable increase in plasticity and cellular heterogeneity^1,2^. The ability of cells to shift states and adopt mixed lineage potential exacerbates during metastasis and associates with poor patient survival^3^. How these atypical differentiated states emerge and drive cancer progression however has remained unclear. Here, we investigate this issue for BRAF-mutant CRC that commonly arise via the serrated pathway^4^ and are linked to poor prognosis via incompletely understood progression steps. Using patient-derived organoids, gene editing and murine tumour models, we show that mutations in RNF43, a regulator of WNT receptor abundance^5^, endow BRAF-mutant CRC with metastatic capacity by driving a non-dividing tumour-intrinsic niche cell (TINC) state that provides growth factor self-sufficiency to the cancer tissue. TINCs are enriched in *RNF43*-mutant, mucinous and metastatic samples of human CRC, and ablation of TINCs lead to re-acquired external growth factor dependency during CRC organoid outgrowth. Together, our findings uncover a mechanism by which tumours leverage cellular plasticity to mediate self-organised stem– and niche-cell interactions. Our results argue that formation of a tumour-intrinsic niche is a prerequisite for BRAF-mutant CRC seeding to distant organs and that interference with niche formation may help avoid metastatic relapse.

## Main

Homeostatic self-renewal of the human colon is controlled by specialised epithelial and mesenchymal niche cells that secrete essential growth factors, including WNT, epidermal growth factor (EGF), Noggin and R-Spondins, to maintain epithelial stem cell populations and guide cellular differentiation toward secretory and absorptive lineages along the crypt-villus axis^6,7^. During cancer development and progression, human CRC cells typically display loss of niche factor-dependency, commonly demonstrated by their capacity to form organoids *in vitro* in absence of growth factor supplementation (Extended Data Fig. 1a). The degree of niche-factor independency of patient-derived or engineered CRC organoids correlates with their metastatic capacity *in vivo*^8,9^.

CRC develops via two mutually exclusive genetic pathways that involve distinct WNT pathway alterations^4,10–12^. In the conventional pathway, initiating mutations in *APC* cause downstream WNT/β-catenin pathway activation which drives a hyperproliferative intestinal stem cell-like growth state that is largely resistant to external signalling cues^1^. By contrast, mutations in *RNF43,* conferring hypersensitivity to WNT^5^, are mainly acquired by BRAF-mutant CRC that develop via the serrated pathway, originating from lesions with flat or sessile morphologies (15-30% of cases)^5,13,14^. *RNF43/BRAF*-mutant CRC display relatively low levels of WNT/β-catenin target gene expression while showing enrichment of differentiated secretory cell types, as reflected by an increased expression of mucin-related genes (Extended Data Fig. 1b)^10,11,15^. Despite displaying low WNT pathway activity, *RNF43/BRAF*-mutant CRC depend on WNT production for their survival^16,17^, exemplified by their sensitivity to Porcupine inhibitors that block WNT secretion (Extended Data Fig. 1c)^9,17^. *RNF43* mutations are absent in earliest *BRAF*-mutant human sessile serrated lesions^11^, and thus must be acquired at later stages of tumour development. The mechanism by which *RNF43* mutations contribute to tumour development and progression has thus far however remained unclear.

### Repairing RNF43 in advanced *BRAF*-mutant CRC eliminates metastatic capacity and niche factor-independency

To address the functional driver role of mutated *RNF43*, we selected HUB040, a microsatellite stable (MSS) *RNF43/BRAF-*mutant human CRC organoid line derived from a primary human mucinous colorectal adenocarcinoma that presented clinically with prominent metastasis^18^. Specifically, the identified RNF43 mutation (P441fs) in this tumour introduces a stop codon downstream of the catalytic RING domain (Fig. 1a), which does not classify as an oncogenic variant^17^, and is therefore predicted to exert (partial) conventional loss-of-function (LOF) effects through deletion of the RNF43 C-terminal tail^19,20^.

**Figure 1.**
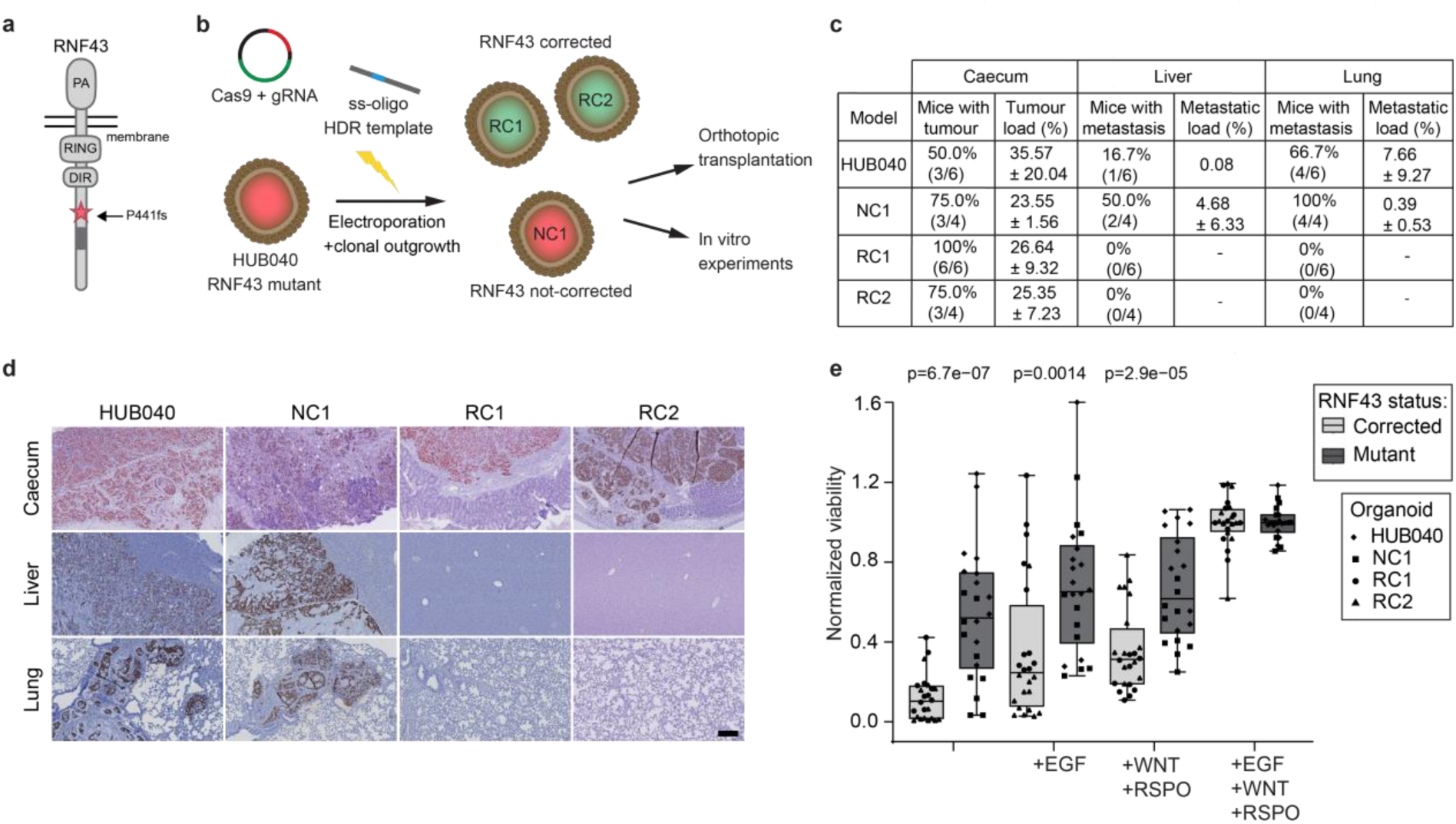
| Repair of *RNF43* mutations in patient-derived organoids blocks metastasis and niche-independent outgrowth. **a.** Schematic of the human RNF43 protein indicating the location of the P441fs mutation. PA: protease associated domain, RING: E3-ligase domain, DIR: Dishevelled-interacting region, dark grey: oncogenic region^19^. **b.** Experimental setup of this study. HUB040 CRC organoids were transfected with a single stranded (ss) HDR template and Cas9-gRNA vector to correct the *RNF43* mutation. Two *RNF43*-corrected clones (RC1, RC2) and non-corrected control clone (NC1) were used for subsequent experiments. **c.** Table showing tumour forming efficiency and tumour/metastatic load in caecum, liver and lung, 120 days after transplantation. Tumour and metastatic load were determined as human nucleoli area over total tissue area. **d.** Representative human nucleoli stainings of caecum, lung and liver of mice, 120 days after transplantation of indicated HUB040 clones. Scale bar = 200 μm **e.** Normalised viability of *RNF43* mutant and *RNF43*-corrected HUB040 clones cultured in CRC medium supplemented with indicated growth factors. Results of six experiments are presented, including one to three replicates per condition (organoid line in a given medium) per experiment (n = 12 per condition in total). Data are normalised against mean value in complete medium per experiment. Kruskal-Wallis test is used to compare *RNF43* mutant and *RNF43*-corrected within each condition.

We employed gene editing to correct the mutated *RNF43* locus to wildtype and obtained two *RNF43*-corrected clones (RC1, RC2) and one non-corrected control clone (NC1) (Fig. 1b and Supplementary Fig. 1a). Whole genome sequencing of all clones confirmed preservation of pre-existing HUB040 driver mutations other than *RNF43* (Supplementary Fig. 1b). Karyotypes of HUB040, NC1 and RC2 were similar, while RC1 was divergent, mainly due to whole genome duplication (Supplementary Fig. 1c-h). To investigate metastatic capacity, we performed orthotopic transplantation of organoids into the caecal wall of immune-deficient mice^21^. After 120 days, HUB040 and NC1 had formed primary tumours that spontaneously metastasized to the lungs and the liver (Fig. 1c,d and Supplementary Fig. 2a). Strikingly, none of the transplanted *RNF43*-corrected organoids induced metastatic lesions (Fig. 1c,d), even though their primary tumour load equalled that of *RNF43*-mutant counterparts (Supplementary Fig. 2a). Thus, by repairing mutant *RNF43* we eliminated the metastatic capacity of these human CRC cells.

To assess whether loss of metastasis is linked to alterations in CRC niche dependency^16^, we performed single cell reconstitution assays in growth factor-depleted conditions. Like HUB040, *RNF43*-mutant NC1 organoids retained their capacity to expand in medium lacking extrinsic growth factors (Fig. 1e and Supplementary Fig. 2b). By contrast, *RNF43*-repaired lines RC1 and RC2 re-acquired a strong dependence on supplementation with all growth factors (WNT, RSPO and EGF), similar to healthy organoids (Fig. 1e, Extended Data Fig. 1a and Supplementary Fig. 2b). These results were unexpected, as the currently known driver mechanism of *RNF43* mutations in human colon organoids explains a state of enhanced WNT sensitivity^13^, but not an alteration in EGF dependency. Our data thus imply a role for *RNF43-P441fs* in promoting cancer cell survival and growth in the absence of growth factor supplementation.

To investigate whether these results are representative for a broader set of human CRC-linked RNF43 mutations, we re-introduced different RNF43 cancer variants in the RC2 (RNF43-corrected) HUB040 background and performed similar organoid reconstitution assays. Introduction of the hotspot mutation *RNF43-G659fs*^22^ and a shorter truncant *RNF43-S323fs*^20^ (Extended Data Fig. 2a) rescued RC2 niche factor-independent growth (Extended Data Fig. 2b), indicating functional similarity with *RNF43-P441fs*. Furthermore, generated HUBU040 *RNF43* knockout clones displayed a retained capacity to grow out in the absence of growth factor supplementation (Extended Data Fig. 2c). We conclude that a broad range of RNF43 mutations facilitate niche-independent growth within this model of advanced *BRAF*-mutant CRC, suggesting a shared mode of action.

### Mutant *RNF43* drives mucinous differentiation and niche factor production

We next aimed to investigate how *RNF43* mutations affect tumour phenotype. Transcriptome comparison of HUB040, NC1 and RC organoid lines uncovered a mutant *RNF43*-driven signature enriched for genes encoding secreted growth factors and mucins (Fig. 2a,b and Supplementary Fig. 3). In line, gene sets representing colon-derived Goblet and Deep Crypt Secretory (DCS) cells were depleted from RC1 and RC2 in comparison to control lines (Fig. 2c)^23,24^. Furthermore, mucus secretion was strongly diminished upon *RNF43* correction, both *in vitro* and *in vivo* (Fig. 2d-f). These findings indicate that *RNF43* mutations facilitate secretory cell differentiation.

**Figure 2.**
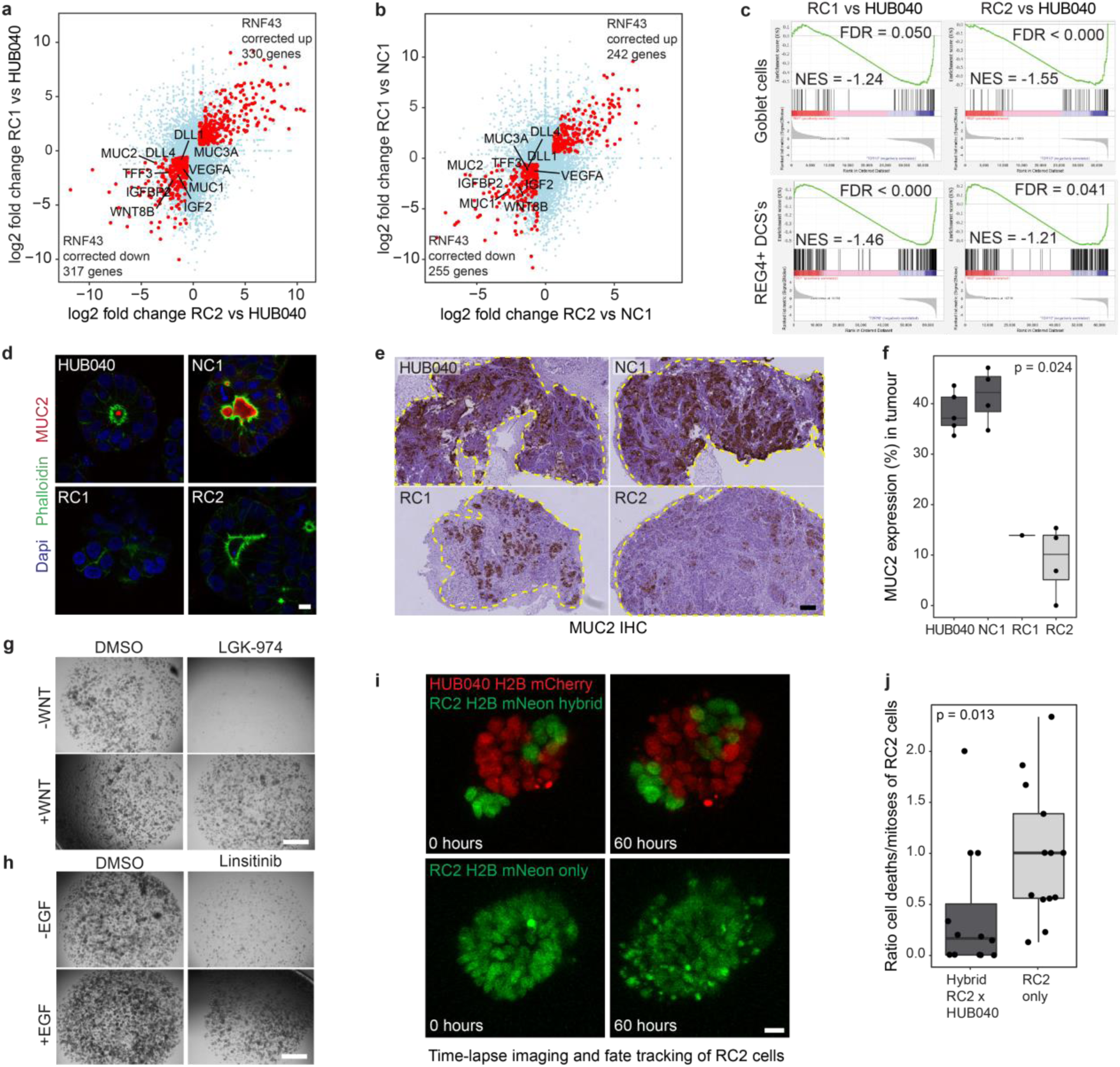
| Mutant *RNF43* drives mucinous differentiation and niche factor production. **a, b.** Scatter plots showing log2 fold change of gene expression between *RNF43*-corrected clones, RC1 (Y-axis) and RC2 (X-axis), and HUB040 (**a**) or non-corrected clone 1 (NC1) (**b**). Genes with significant differential expression (FC > 1.5 and p < 0.05) in both comparisons are indicated as larger red dots. Downregulated genes of interest are annotated. Total number of genes that are up– or downregulated in both comparisons is plotted in the corners. **c.** GSEA plots showing enrichment of a colonic goblet cell signature^24^ and REG4+ deep crypt secretory cell (DCS) signature^23^ in HUB040 compared to RC1 or RC2. **d.** Representative confocal microscopy images of organoids stained for mucin2 (MUC2, in red), actin (phalloidin, green) and nuclei (dapi, blue). Scale bar = 10 µm. **e.** Representative IHC images for mucin2 (MUC2) on primary subcutaneous tumours. Scale bar = 250 µm. **f.** Box plots showing the percentage of MUC2+ tumour area in primary subcutaneous tumours. Significance was evaluated using a Krusal-Wallis test. **g, h.** Representative low-magnification brightfield images of HUB040 organoids treated with 100 nM PORCN inhibitor LGK-974 (**g**), 1 µM IR/IGFR inhibitor Linsitinib (**h**) or DMSO in CRC medium with or without WNT surrogate (**g**) or EGF (**h**) supplementation for 14 days, passaged after 7 days. Scale bar = 1000 μm. **i.** Representative 3D projected images of time-lapse series of hybrid HUB040-RC2 or only RC2 organoids in CRC medium at the start (0 hours) and end (60 hours) of the time-lapse; nuclei are visualized by expression of H2B-mNeon (RC2) or H2B-mCherry (HUB040). Scale bar = 20 µm. **j.** Analysis of RC2-cell fate over time in the two conditions described in (**i**). Cell death and mitotic events were counted for all RC2-H2B-mNeon cells in 13-14 movies per condition. Graph depicts the ratio of the number of cell death over mitotic events.

Notably, several known niche factors, including *IGF2*, *VEGFA*, *DLL1*/*4* and *WNT8B* were present in the *RNF43*-mutant signature (Fig. 2a,b and Supplementary Fig. 3). Moreover, blocking upstream WNT or IGF signalling diminished HUB040 organoid reconstitution capacity, which was rescued by respectively WNT ligand or EGF supplementation (Fig. 2g,h). To examine whether *RNF43* mutations promote the generation of self-produced niche-factors, essential for tumour survival and growth, we generated hybrid co-cultures of *RNF43*-mutant and *RNF43*-corrected organoids and monitored cellular death and division events in growth factor-depleted medium (Fig. 2i and Supplementary Videos 1,2)^25^. While cultures of only *RNF43*-corrected organoids displayed a high cell death/mitosis ratio, this ratio was reversed when they shared a lumen with *RNF43*-mutant cells (Fig. 2j). These results thus indicate that HUB040 CRC survival relies on paracrine signalling facilitated by mutant RNF43.

### Mutant *RNF43* promotes tumour-intrinsic niche cell formation

Notably, our earlier work showed that inactivation of *Rnf43* and its paralogue *Znrf3* in the mouse small intestine induces hyperplasia of stem cells as well as secretory Paneth cell populations that function as an intra-epithelial stem cell niche^5,26^. We thus hypothesized that *RNF43* mutations promote the formation of specialized niche factor-producing cells within human CRC. To assess this issue, we examined single cell HUB040 profiles that spanned seven clusters with distinct expression patterns as shown by unsupervised clustering (Fig. 3a and Extended Data Fig. 3a). Noticeably, one cell population (cluster 1) entirely lacked expression of proliferation and stem cell-related genes^27^ (Fig. 3b,c and Extended Data Fig. 3b). Instead, this subset displayed prominent hallmarks of differentiated secretory niche cells, exemplified by gene sets related to Goblet cells (Fig. 3d), Deep Crypt Secretory (DCS) cells (Fig. 3e)^23,24^, glycolysis and hypoxia (Extended Data Fig. 3c-e)^28,29^, mTOR activation and the unfolded protein response (Extended Data Fig. 3e)^30–32^. Hence, we identify cluster 1 as a non-proliferating, niche factor-producing secretory cancer cell subset that we call tumour-intrinsic niche cells (TINCs).

**Figure 3.**
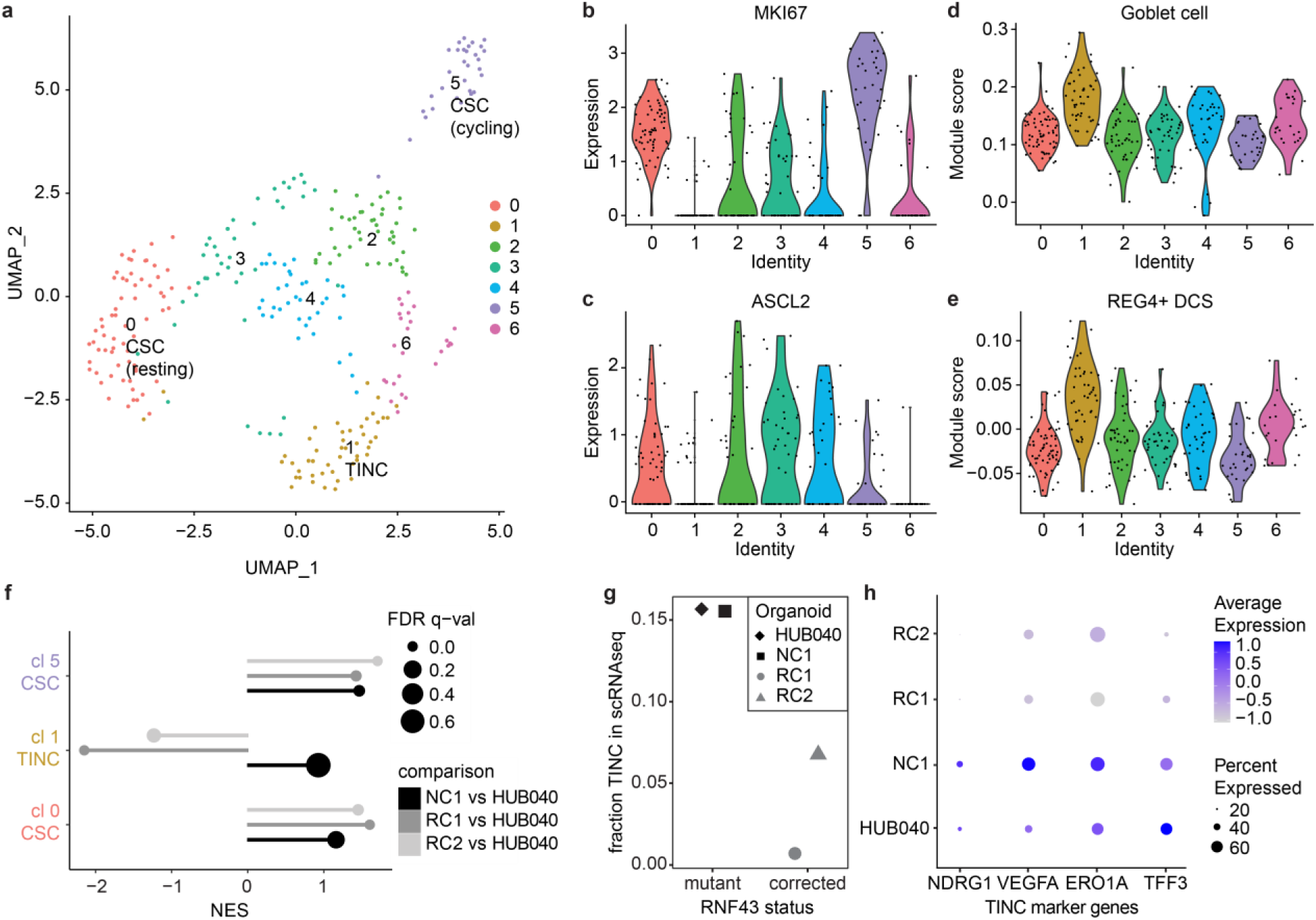
| Mutant *RNF43* promotes tumour-intrinsic niche cell formation. **a.** Reduced dimensionality (UMAP) visualisation of single cell RNA sequencing data of HUB040 organoids cultured in CRC medium. Results of unsupervised clustering are colour-coded per cluster. **b, c.** Violin plots showing scaled single cell expression levels of proliferation marker MKI67 (**b**) and cancer stem-cell marker ASCL2 (**c**) grouped by cluster. **d, e.** Violin plots showing single cell module scores for expression of genes from a colonic goblet cell signature^24^ (**d**) and REG4+ deep crypt secretory cell (DCS) signature^23^ (**e**) grouped by cluster. **f.** GSEA comparing *RNF43*-corrected and non-corrected clones with HUB040 naïve by marker gene signatures for three HUB040-specific clusters (CSC clusters 0 and 5 and TINC cluster 1). x-axis shows the normalised enrichment score (NES) and size of the dots represents the false discovery rate (FDR). **g.** Plot showing the fraction of cells that were scored as TINC in the analysis of single cell RNA sequencing data of the four indicated organoid lines. **h.** Dot-plot showing the average expression and fraction of cells expressing indicated TINC marker genes.

Of note, the TINC gene signature was significantly downregulated in both RNF43-corrected clones (Fig. 3f). scRNAseq analysis of all clones in complete medium revealed that the fraction (16%) of HUB040 and NC1 cells that displayed high TINC signature expression dropped to 1-7% of cells within RNF43-corrected clones (Fig. 3g). In agreement, expression of several prominent TINC markers were diminished in RNF43-corrected organoids (Fig. 3h and Extended Data Fig. 3f). Based on these findings, we conclude that TINC formation is promoted by mutant RNF43. Strikingly, TINCs shared transcriptional identity with previously identified high-relapse cells (HRCs) in CRC, a high-plasticity cell state (HPCS) in lung cancer and a basal-like PDAC signature (Extended Data Fig. 3g-i)^33–35^, suggesting that TINCs combine their niche role with an increased capacity to metastasize to distant organs.

### *RNF43*-mutant organoids self-organise into proliferating stem cells and non-proliferating supporting niche cells

To perform an in-depth characterisation of TINC cells, we introduced an IRES-linked mNeon green fluorophore into the *NDRG1* gene that prominently marks TINCs in HUB040 (Fig. 4a,b). Non-dividing NDRG1^high^ TINC reporter cells were heterogeneously distributed in HUB040 organoids interspersed with proliferative NDRG1^low^ cells as shown by KI67 labelling (Fig. 4c) and EdU incorporation (Fig. 4d). Despite their differentiated and non-proliferative appearance, TINCs remained highly plastic, as isolated single NDRG1^high^ cells gave rise to fully grown organoids in which a heterogeneous distribution of TINCs was restored (Fig. 4e,f). Thus, *RNF43/BRAF*-mutant HUB040 CRC organoids display phenotypic heterogeneity and are capable of self-organising into a mixture of non-proliferative TINCs and stem-like cell populations.

**Figure 4.**
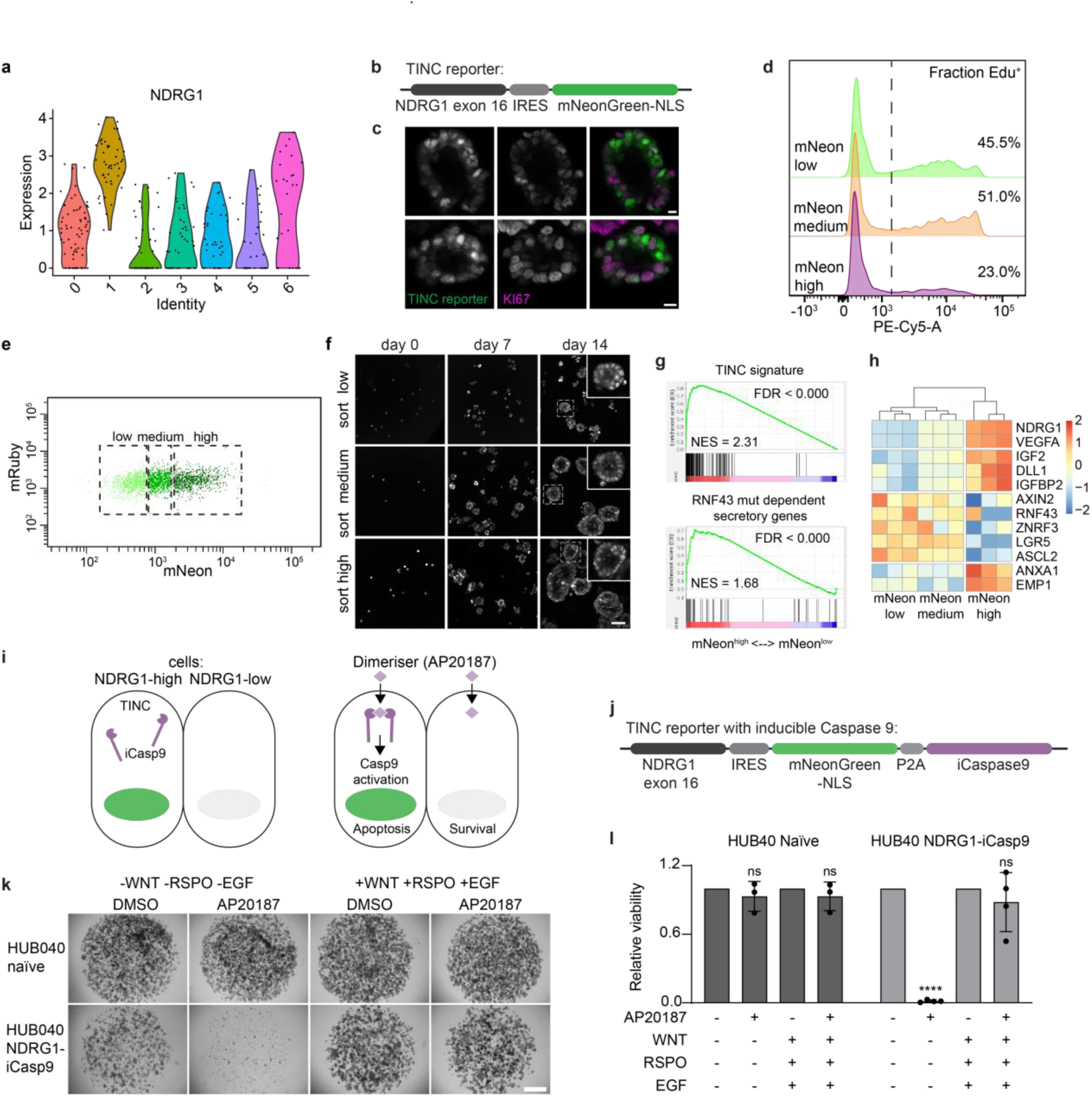
| *RNF43*-mutant organoids self-organise into proliferating and supporting cells that are essential for niche-independent survival. **a**. Violin plot showing single cell expression levels for NDRG1 grouped by HUB040 cluster. **b.** Schematic of the endogenous TINC reporter construct. **c.** Representative confocal microscopy pictures of TINC reporter-positive cells (mNeon) in HUB040 organoids, combined with KI67 immunostaining. Scale bar = 10 µm. **d.** FACS plots of HUB040 TINC reporter organoids gated for low, medium or high mNeon signal showing the result of EdU labelling after 16h incubation with EdU. **e.** FACS plot of mNeon signal for TINC reporter organoids: sorted populations are indicated. A constitutive mRuby signal was present in the reporter for selection. **f.** Representative confocal microscopy images of HUB040 TINC reporter organoids 0, 7 and 14 days after sorting. Scale bar = 50 µm. **g**. GSEA plots showing enrichment of the TINC signature that was derived from single cell RNAseq and the ‘RNF43-mutation dependent secretory gene signature’ as defined in Supplementary Figure 3 for TINC reporter high versus low cells. **h.** Heatmap showing Z-scores for the expression of selected genes (TINC markers, growth factors, WNT target genes and regenerative genes) in the TINC reporter low, medium and high cells. **i.** Cartoon depicting the mechanism of the inducible Caspase 9 (iCasp9) construct. **j.** Schematic of the endogenous iCaspase TINC reporter construct. **k.** Representative low-magnification brightfield images of a single cell reconstitution assay of HUB040 naïve and NDRG1-iCasp9 organoids treated with DMSO or 10 nM AP20187 for 14 days. Scale bar = 1000 μm. **l.** Quantification of reconstitution assays shown in (**k**) using a CellTiter-Glo assay on day 14. Graph showing average levels of luminescence normalised to DMSO control of n=3-4 experiments +/− S.D. One-way ANOVA.ns, not significant; **** (P < 0.0001).

Examination of the transcriptomes of isolated NDRG1^high^ compared to NDRG1^low/medium^ cells robustly confirmed enrichment of hallmark gene sets for TINCs (Extended Data Fig. 4a) and high relapse cells (HRC; Extended Data Fig. 4b)^33^ as well as signature genes (Fig. 4g), including marker genes of regeneration (*ANXA1*) and metastatic relapse (*EMP1*) (Fig. 4h)^33,36^. Importantly, and in line with our model, the majority of secretory genes linked with mutant RNF43 expression (Supplementary Fig. 3) were enriched in NDRG1^high^ cells (Fig. 4g and Extended Data Fig. 4c) including the bona fide niche factors *IGF2, Dll1, IGFBP2* and *VEGFA* (Fig. 4h). Intriguingly, TINCs displayed a relative decrease in WNT target gene and stem cell marker expression, including *RNF43* itself (Fig. 4h). These results support a model in which *RNF43* mutant-tumours are capable of forming an intra-tumoural WNT gradient along which lineage diversification may occur, even though all cells are *RNF43*-mutant. We conclude that *RNF43* mutations promote the differentiation of a fraction of tumour cells into a niche factor-producing state to ensure self-sustained growth of these tumours.

### Adjustable WNT signalling balances the formation of stem cells and supporting tumour niche cells

We next aimed to address the relationship between WNT signalling, TINC formation and tumour growth. In human CRC, *BRAF/RNF43* mutant tumours are mutually exclusive with *APC* mutations, that drive high and incessant WNT/β-catenin signalling activation^12^. To assess how high WNT levels affect TINC formation, we truncated *APC* within HUB040 organoids. Compared to HUB040^APC+/−^ clones, HUB040^APC-/-^ organoid clones displayed prominent WNT/β-catenin target gene expression, enhanced nuclear β-catenin levels and an acquired resistance to treatment with WNT secretion inhibitors (Extended Data Fig. 5a,b and Supplementary Fig. 4a-c). In line with earlier work^37^, homozygous APC mutation are not compatible with efficient growth of serrated CRC cells, as HUB040^APC-/-^ organoids were outcompeted by HUB040^APCwt^ organoids during co-culture (Supplementary Fig. 4d). Furthermore, mutant APC reduced TINC signature expression, while enriching for transit amplifying (TA) and stem-like cell populations (Extended Data Fig. 5c,d)^38^. Single cell analysis confirmed that mutant APC mediated a shift in HUB040 cell identity and composition, driving an increased fraction of WNT-high stem-like cancer cells and a reduced fraction of TINC signature-expressing cells (Extended Data Fig. 5e,f and Supplementary Fig. 4e-g). We thus conclude that mutant *APC* interferes with intra-epithelial niche formation in HUB040 that is otherwise facilitated by mutant *RNF43*.

Based on these results, we hypothesized that adjustable levels of WNT signalling are vital for *BRAF/RNF43*-mutant tumour self-organising capacity. HUB040 organoids treated with WNT inhibitors for 24 and 48 hours (Supplementary Fig. 4h-j) revealed an incremental downregulation of proliferative, stem and TA signatures and an increase in differentiation signatures, including TINC (Extended Data Fig. 5c,d). In line with these findings, the number and brightness of TINC reporter cells in HUB040 increased upon WNT inhibitor treatment (Extended Data Fig. 5g and Supplementary Fig. 4k). Thus, although *BRAF/RNF43*-mutant mucinous cancers are wired towards lower levels of WNT signalling that are optimal to facilitate secretory differentiation, a threshold of WNT signalling in a subset of cancer cells is still required to prevent terminal differentiation and maintain self-renewal of proliferative cells.

### Ablation of TINCs diminishes niche-independent reconstitution in RNF43-mutant CRC

To investigate how HUB040 CRC organoid survival and growth functionally depends on the generation of TINCs, we introduced an inducible caspase-9 cassette downstream of the *NDRG1* locus (Fig. 4i,j)^39^. Treatment with the dimerizing compound AP20187 induced specific ablation of NDRG1-high TINC cells (Supplementary Video 3), which fully abolished the capacity of HUB040 to reconstitute organoid outgrowth in growth factor-depleted medium (Fig. 4k,l). Strikingly, the organoid outgrowth in TINC-depleting conditions was fully rescued by extrinsic supplementation with EGF, WNT and RSPO (Fig. 4k,l). We conclude that cyto-ablation of TINCs phenocopies RNF43-correction in HUB040 organoids, strongly supporting a model in which mutant RNF43-driven formation of a tumour-intrinsic niche contributes to growth factor self-sufficiency and an increased metastatic capacity of this subset of CRC.

### *Rnf43* mutations drive TINC formation and metastasis of *Braf*-mutant intestinal tumours ***in vivo***

Combined, our data suggest that the introduction of RNF43 mutations drive progression of BRAF-mutant tumours by promoting the formation of an intra-tumoural niche. While *RNF43* mutations are scarce in early lesions of BRAF-mutant CRC^11^, they accumulate to 60% of cases in advanced stages of disease^11^. To examine whether loss of *RNF43* accelerates CRC progression *in vivo*, we utilised a tamoxifen-inducible VillinCre^ER^ transgene, specific to the cells of the intestinal epithelium, to generate Villin-Cre-ERT2 *Braf^V600E^ Trp53^fl/fl^* (BP), Villin-Cre-ERT2 *Braf^V600E^ Rnf43^fl/f^* (BRNF) and Villin-Cre-ERT2 *Braf^V600E^ Trp53^fl/fl^ Rnf43^fl/fl^* (BPRNF) mouse models (Fig. 5a). Within the three-allele hit BPRNF model, tumourigenesis was strongly accelerated compared to BP and BRNF (Fig. 5b). Strikingly, while metastasis was absent in BP mice, BPRNF mice displayed a high incidence of metastasis (71%), with prominent dissemination to mesenteric lymph-nodes (mLN) and liver (Fig. 5c,d). Furthermore, BPRNF-tumours displayed a mucinous (Muc5ac-high) phenotype (Fig. 5e) and were transcriptionally enriched for genes encoding for both mucins and niche factors, including WNTs (Fig. 5f). Finally, single cell RNA analysis revealed that TINC-high clusters (clusters 5, 7 and 10) were enriched in BPRNF tumours (Fig. 5g-i). Together, these data support a model in which inactivating mutations in *RNF43* accelerate CRC progression *in vivo* via formation of a tumour-intrinsic niche that endows cells with an increased metastatic capacity.

**Figure 5.**
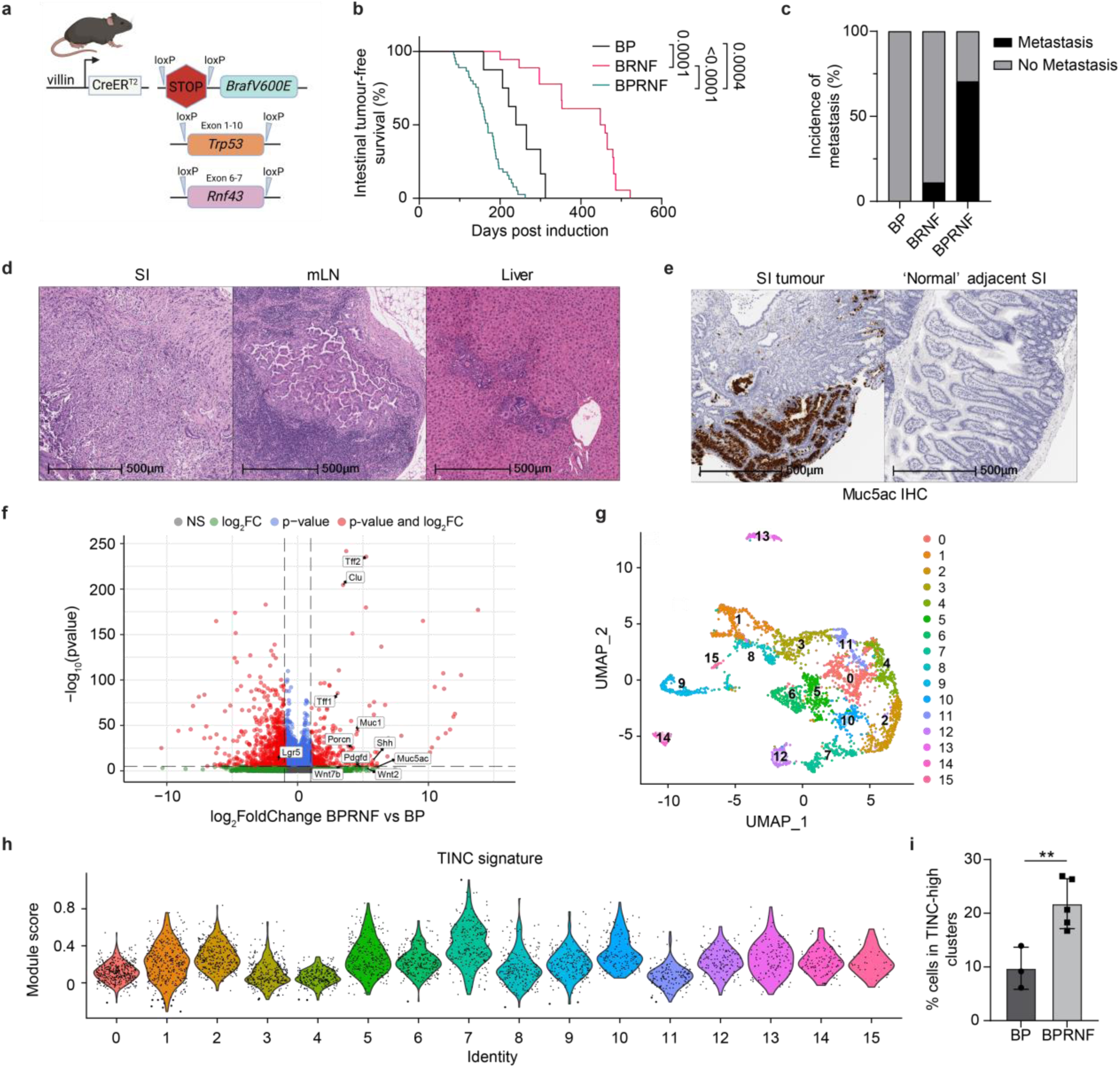
| Rnf43 loss drives metastasis and TINC formation in vivo. **a.** Schematic overview of the genetic crossing strategies to generate: villin-cre-ERT2 *Braf^V600E/+^ Trp53^fl/fl^* (BP), villin-cre-ERT2 *Braf^V600E/+^ Rnf43^fl/fl^* (BRNF) and villin-cre-ERT2 *Braf^V600E/+^ Trp53^fl/fl^ Rnf43^fl/fl^* (BPRNF) mouse models. Cre, Cre-recombinase; ER, Oestrogen receptor; LoxP, Cre-Lox recombination site. **b.** Kaplan-Meier survival analysis of intestinal tumour-free survival in Braf-driven models of colorectal cancer with or without additional *Trp53* and or *Rnf43* loss. Log-rank (Mantel-Cox) tests for significance. Tumour-free mice (indicated by vertical ticks on the survival curve) were censored due to health reasons not related to intestinal tumourigenesis. Median overall survival presented as days post tamoxifen induction (DPI): BP 252 DPI (n = 9; 4M, 5FM), BRNF 454 DPI (n = 18; 7M, 11FM) and BPRNF 169 DPI (n = 46; 24M, 22FM). **c.** Percent incidence of BP, BRNF and BPRNF mice with metastasis. Tumour-free censored mice free of intestinal tumourigenesis were removed from metastatic analysis. BP n = 0/7, BRNF n = 2/18 (0M 2FM) and BPRNF n = 31/44 (14M, 17FM). **d.** Representative H&E images of BPRNF primary small intestinal (SI) tumour, mesenteric lymph node (mLN) metastasis and liver metastasis. Scale bars 500 μm. **e.** Representative image of *Muc5ac* IHC in the villin-cre-ERT2 *Braf^V600E/+^ Trp53^fl/fl^ Rnf43fl/fl* (BPRNF) mouse model at sampling comparing SI tumour and normal adjacent SI tissue. Scale bar 500 μm. **f.** Volcano plot showing the differential gene expression within epithelial cells of BPRNF versus BP tumours, indicating significant genes of interest. ns = not significant. **g.** UMAP visualization of single cell RNA sequencing data of isolated epithelial cells from 3 BP and 5 BPRNF tumours. Clustering results are colour-coded per cluster. **h.** Violin plot showing single cell module scores for expression of genes from the TINC signature grouped by cluster. **i.** Percentage of cells mapping to the TINC-high clusters (5, 7 and 10) for individual BP and BPRNF tumours. Unpaired t-test for significance. ** (P < 0.01).

### TINCs are enriched in *RNF43*-mutant human CRC and metastatic lesions

We next explored the broader relevance of TINCs for human CRC. TINC marker gene expression was enriched in *RNF43*-mutant and mucinous CRC cases of the cancer genomics atlas (TCGA) (Extended Data Fig. 6a,b). Furthermore, when classifying tumours based on CRC consensus molecular subtype (CMS)^40^ TINC signatures were lowest in the WNT-high subtype CMS2 that is enriched for APC-mutant tumours (Extended Data Fig. 6c). These findings were corroborated by analysis of 18.320 single cell transcriptomes derived from 28 human CRC tumours, of which four were *RNF43* mutant^10^ (Extended Data Fig. 6d). Supervised clustering with batch correction resolved eight shared CRC cell clusters (Extended Data Fig. 6e). In comparison with *RNF43*-wildtype CRC, the *RNF43*-mutant CRC subset displayed enrichment for cells in cluster 3 and 5 (Extended Data Fig. 6f) that exhibit goblet cell (cluster 5) or TINC markers (cluster 3), including *NDRG1* (Extended Data Fig. 6g,h). Thus, *RNF43*-mutant CRC subsets are not only enriched for TINC marker expression but also display an overall increased fraction of TINCs in comparison to RNF43-wildtype CRC. Additionally, TINC gene expression was enhanced in lung and liver CRC metastatic samples compared to primary tumours (Extended Data Fig. 6i), indicating that enhanced TINC profiles associate with advanced disease.

## Discussion

The heterogeneous identities of cancer cells are shaped by both cell-intrinsic mutations and cell-extrinsic signalling cues^1^. Here, we uncover a genetic cancer driver event that enhances the plasticity of cancer cells by promoting their entry into a non-canonical, secretory niche cell state. We show that this increase in phenotypic plasticity endows cancer cells with the capacity to polarize into stem and niche cell populations that functionally interact, conferring a state of growth signalling autonomy to the cancer epithelium. Importantly, these self-organizing properties are linked to an increased capacity of cancer cells to metastasize to distant organs. Due to the overlap in identity of TINCs and HRCs^33^, we propose a model in which the seeding of niche-like cells precedes the formation of CSC populations in distant organ metastasis. This model is reminiscent of the process of intestinal regeneration and repair, in which symmetry breaking events of a uniform population of fetal-like epithelial cells first lead to the formation of niche cells after which Lgr5-high stem cells appear^41^. At the same time, our findings provide a mechanistic explanation for the phenomenon of ‘histostasis’, in which metastasized cancer tissues display an inherent ability to self-organise and recreate a morphology similar to the primary tumour, albeit growing in a different organ microenvironment^42^. The capability of forming interdependent stem and niche cell states likely extends to other cancer types, since a specialized Wnt-producing niche was identified in lung adenocarcinoma^43^. Intriguingly, and in line with our results, the acquirement of growth signalling autonomy of circulating breast cancer cells was shown to improve their fitness and metastatic potential^44^. Taken together, a comprehensive understanding of the pathways and mechanisms driving tumour niche self-organization will not only provide novel vulnerabilities but also reveal targets that may be harnessed to prevent disease progression and metastatic recurrence.

## Methods

### Patient material

The collection of colorectal tissue for the generation of healthy colon and CRC organoids was performed according to the guidelines of the European Network of Research Ethics Committees (EUREC) following European, national and local law. In all cases, patients signed informed consent after ethical committees approved the study protocols. Organoids identified by HUB codes *HUB-02-B2-040* (HUB040, CRC) and *HUB-02-A2-040* (HUB040N, normal colon) are catalogued at https://huborganoids.nl/ and can be requested at. Distribution to third (academic or commercial) parties will have to be authorized by the Biobank Research Ethics Committee of the University Medical Center Utrecht (TCBio) at request of Hubrecht Organoid Technology (HUB). P26T (CRC) was established in a previous study^17^.

### Organoid culture

Healthy colon organoids and CRC organoids were cultured according to previous protocols^17^. In short, organoids were cultured in drops of Matrigel (Corning) or BME (R&D systems), and the medium was refreshed every two or three days. CRC organoids were passaged using TrypLE Express (Life Technologies) incubation at 37 °C. Healthy colon organoids were passaged using mechanical shearing (pipetting). Healthy colon organoids were cultured in advanced DMEM/F12 medium (Invitrogen), supplemented with 1% Penicillin/Streptomycin (P/S, Lonza), 1% Hepes buffer (Invitrogen), 1% Glutamax (Invitrogen), 1x B27 (Invitrogen), 10 mM Nicotinamide (Sigma-Aldrich), 1.25 mM N-acetylcysteine (Sigma-Aldrich), 50 ng/mL EGF (PeproTech), 500 nM TGF-b type I receptor inhibitor A83-01 (Tocris), 10 µM P38 inhibitor SB202190 (Sigma-Aldrich), 50% Wnt3a-CM (homemade) or 1% surrogate WNT-CM (homemade)^45^, 10% Noggin-CM (homemade), 20% Rspo1-CM (homemade) and 100 µg/mL Primocin (Invitrogen) (full HCO medium). Colorectal carcinoma organoids were cultured in medium without EGF, WNT and RSPO (CRC medium), unless stated otherwise. *RNF43*-corrected (RC), non-corrected (NC) HUB040 clones were maintained in full HCO medium to prevent negative selection against growth factor dependence.

### CRISPR strategy and design

To correct the *RNF43* P441fs mutation back to wildtype, HUB040 organoids were electroporated with a single stranded HDR template (ss-oligo) and Cas9-gRNA. The ss-oligo was designed according to previously established optimal rules.^46,47^ Briefly, the *RNF43* mutation in HUB040 is a five base pair duplication, which was corrected in the template. The template was an asymmetric template oligo complementary to the non-target strand overlapping the Cas9 cut site with 36 bp extension on the PAM-distal side, and with a 91-bp extension on the PAM-proximal side of the break. The ss-oligo was ordered at Integrated DNA Technologies with phosphorothioate modifications of the last two nucleotide bonds at the 3’ and 5’ end (sequence: TATGCACAAGGCTGGGGAATGAGCCACCTCCAATCCACCTCACAGCACCCTGCTGCTTGCCCAGTGCCCCTACGCCGGGCCAGGCCTCCAGACAGCAGTGGATCTGGAGAAAGCTATTGCACAGAACGC). A gRNA was chosen using the Benchling tool (https://benchling.com) that created a double stranded break close to the mutation (sequence: CCAGATCCACTGCTGTCAGG) and cloned in pSpCas9(BB)-2A-Puro (Addgene, 48139).

The following gRNAs were used to introduce RNF43 mutations: (RNF43 S322fs: GATTCATTTTCCCAGTCCCT and RNF43 G659fs: CCACAGAGGAAAAGGCGGGG). and RNF43 KO-exon3: ACCCGCTGTACCTGTGCAATG) APC mutations were introduced by targeting the mutation cluster region (MCR) in the *APC* gene using previously reported gRNA (sequence: TACATCTGCTAAACATGAG).^48^

Reporter targeting vector was generated by altering *HR180-LGR5-iCT* plasmid (Addgene #129094) into a plasmid compatible with insertion of homology arms using golden-gate cloning^49^. pase-T2A-tdTomato sequence was removed and swapped for mNeonGreen-NLS using a combination of Q5® High-Fidelity DNA Polymerase and the In-Fusion HD Cloning kit (Takara). Final targeting vector was generated by insertion of homology arms (∼600bp each) flanking the targeted region downstream of NDRG1 to obtain the NDRG1-specific *IRES-mNeonGreen-NLS-WPRE-bGHpA-EF1-Ruby-T2A-Puro-SV40pA* targeting plasmid. The same homology arms were inserted into an *IRES-mNeonGreen-NLS-P2A-iCaspase9-WPRE-loxP-PGK-PuroRPuro-SV40pA-loxP* targeting plasmid. gRNAs were cloned into pSpCas9(BB)-2A-Puro using golden gate cloning (sequence: CCGGACTCTGATCTCTGTAG).

### Electroporation

Organoid electroporation was performed as previously described^50^. Briefly, organoids were grown in antibiotic-free medium including 10 µM Y-27632 (Selleck chemicals) for two days before electroporation. 24 hours before electroporation, 1.25% DMSO was added to the culture medium. For electroporation, the NEPA21 electroporator was used with the configuration reported by Fuji et al.^50^. For *RNF43* correction and generation of TINC-reporter, 5 μg of Cas9-gRNA and 5 μg template were transfected. For the generation of the iCaspase reporter, 4 μg of Cas9-gRNA and 11 μg template were transfected and voltage was adjusted to 190V. For other CRISPR targeting, 10 μg of Cas9-gRNA was transfected. Organoids were recovered for two days by adding medium supplemented with 10 µM Y-27632. From five days post-electroporation organoids were grown in full medium. For CRISPR-engineered organoids, single clones were established by manual picking of individual organoids derived from single cells and genotyped. Reporter lines were selected by puromycin treatment and sorted for the constitutive mRuby fluorophore in case of the NDRG1-mNeon reporter. Single cell derived clones were picked manually and genotyped.

### Genotyping

For genotyping of genome-edited organoids, genomic DNA was isolated using QIAamp DNA micro kit (Qiagen). PCR amplification performed using GoTaq Flexi DNA polymerase (Promega) and products were loaded on an agarose gel. Gel fragments of expected sizes were isolated using QIAquick gel extraction kit (Qiagen) and sequenced by Sanger sequencing. Knockout and Knockin efficiencies were determined using the TIDE^51^ (http://shinyapps.datacurators.nl/tide/) and TIDER^52^ (http://shinyapps.datacurators.nl/tider/) tools respectively. Manual verification of genotypes for clonal lines was done in Benchling.

HUB040 NDRG1-IRES-mNeon and iCaspase-NDRG1-mNeon clonal organoid lines were genotyped to confirm homozygous integration of the knock-in construct using gel electrophoresis of PCR products.

Primers used for genotyping are listed in Supplementary Table 1.

### Whole-Genome Sequencing and mapping

Genomic DNA of naïve HUB040 and clonal organoid lines was isolated using QIAamp DNA micro kit (Qiagen) and DNA libraries for Illumina sequencing were generated by using standard protocols (Illumina) Libraries were sequenced on a NovaSeq 6000 S4 platform with 2 x 150 bp read length to 30 times base coverage.

Sequencing reads were mapped against human reference genome GRCh38 by using Burrows-Wheeler Aligner v0.7.17 mapping tool^53^ with settings ‘bwa mem –c 100 –M’. Sequence reads were marked for duplicates by using Sambamba^54^ v0.6.8 and sequence read quality scores were recalibrated with GATK^55^ BaseRecalibrator v2.7.2. CNVs detection was performed with Control-FREEC^56^ v11.56. Full pipeline description and settings are also available at: https://github.com/UMCUGenetics/IAP.

### Mutation calling and filtering

Mutation calling and filtering was performed on multi-sample vcfs generated by IAP using HaplotypeCaller from GATK v.4.1.3.0. GATK’s VariantFiltration was used for variant quality evaluation with options: “––filter-expression ‘QD < 2.0’ ––filter-expression ‘MQ < 40.0’ ––filter-expression ‘FS > 60.0’ ––filter-expression ‘HaplotypeScore > 13.0’ ––filter-expression ‘MQRankSum < –12.5’ ––filter-expression ‘ReadPosRankSum < –8.0’ ––filter-expression ‘MQ0 >= 4 && ((MQ0 / (1.0 * DP)) > 0.1)’ ––filter-expression ‘DP < 5’ ––filter-expression ‘QUAL < 30’ ––filter-expression ‘QUAL >= 30.0 && QUAL < 50.0’ ––filter-expression ‘SOR > 4.0’ –– filter-name ‘SNP_LowQualityDepth’ ––filter-name ‘SNP_MappingQuality’ ––filter-name ‘SNP_StrandBias’ ––filter-name ‘SNP_HaplotypeScoreHigh’ ––filter-name ‘SNP_MQRankSumLow’ ––filter-name ‘SNP_ReadPosRankSumLow’ ––filter-name ‘SNP_HardToValidate’ ––filter-name ‘SNP_LowCoverage’ ––filter-name ‘SNP_VeryLowQual’ ––filter-name ‘SNP_LowQual’ ––filter-name ‘SNP_SOR’ –cluster 3 –window 10”.

Subsequently, SNPEffFilter^57^, SNPSiftDbnsfp (database dbNSFP3.2a^58^;, GATK VariantAnnotator (database COSMIC v.89), and SNPSiftAnnotate (database GoNL release 5) were used for variant annotation.

Finally, to obtain catalogues of high-quality somatic mutation calls, we applied post-processing filtering steps, per sample, as described below (all scripts are available at: https://github.com/ToolsVanBox/SMuRF).

Briefly, only variants were considered if present on autosomal chromosomes, passed VariantFiltration with a GATK phred-scaled quality score ≥ 100, had a base coverage of at least 10X, had a mapping quality (MQ) score of 60, did no overlap with single nucleotide polymorphisms (SNPs) in the Single Nucleotide Polymorphism Database v146 and a panel of unmatched normal human MSC and fetal genomes (BED-file available upon request), had a GATK genotype score (GQ) of 99 (indel/sbs in clonal sample, or indel in paired control) or higher than 10 (sbs in paired control), had a variant allele frequency (VAF) of >=0.1 to exclude *in vitro* accumulated mutations.

### Identification of driver mutations

Single base substitutions or indels were considered as driver mutations when they have a MQ of 30, a minimal base coverage of 10x, had a VAF equal or higher than 0.1, or present in driver genes COSMIC cancer gene consensus (version of 9/5/2019), were also annotated as missense, frameshift, stop-gain, insertion or deletion, had either a high or moderate expected effect as annotated by SnpEff, and were not present in the Single Nucleotide Polymorphism Database v146 and a panel of unmatched normal human MSC and fetal genomes (BED-file available upon request). Oncoplot was generated with the ComplexHeatmap^59^ v2.12.0 package from Bioconductor.

Structural variant and chromosomal copy-number alteration calling was performed using the GRIDSS-purple-linx^60^ pipeline developed at the Hartwig Medical Foundation.

### *In vivo* subcutaneous tumourigenesis

Animal experiments were approved by the Competent Authority, The Netherlands (License number AVD115002016614), which is advised by the Animal Ethics Committee. Animal work protocol (614-1-22) was approved by the Animal Welfare Body and were performed in accordance with the Dutch Law on Animal Experiments and the European Directive 2010/63/EU. Healthy 8–10 week old 25–30 g male *NOD.Cg-Prkdc^scid^ Il2rg^tm1Wjl^/SzJ* (NSG) mice were supplied by Charles River. Animals were randomly allocated in groups of three mice into individually ventilated cages. Animals were kept at room temperature under 12 hours light/dark cycles and received standard chow pellets and water ad libitum. HUB040 organoids were dissociated five days after passaging into single-cells by TrypLE Express incubation for 5 min at 37 °C. A cell-matrigel (1:1) suspension was prepared with 5 × 10^6^ cells/mL. Using a 1 mL syringe and 25G 0.5 × 16 mm needles, 100 µl of the suspension were subcutaneously injected into the right flank of animals. Animal welfare was monitored by physical appearance, behaviour, and body weight. Animals were sacrificed when the tumour volume (V) reached 1.5 mm^3^, as measured by V = 1/2 × (smaller diameter^2^ × larger diameter). Tumour tissue was harvested for further immunohistochemical analyses. Sample sizes are outlined in the Figure legends. Mice were randomly assigned to experimental groups. No blinding method was used for injection. No animals were excluded from downstream analysis.

### Orthotopic implantation in the caecum of NSG mice

In order to evaluate the tumourigenic capacity of the organoids, murine orthotopic caecum-implantation model^21^ was used as described in animal work protocol (614-1-22). In summary, day before implantation, HUB040 organoids were dissociated into single cells and 2.5 × 10^5^ cells were plated in 10 μL drops of neutralized Rat Tail High Concentrated Type I Collagen (Corning, C3867). Cultures were allowed to recover overnight at 37 °C, 5% (vol/vol) CO2. NSG mice were treated with a s.c. dose of Carprofen (5 mg/kg, Rimadyl^TM^) 30 min before surgery and were subsequently sedated by using isoflurane inhalation anaesthesia [∼2% (vol/vol) isoflurane/O2 mixture]. The caecum was exteriorized through a midline abdominal incision and a single collagen drop containing organoids was surgically transplanted in the cecal submucosa. Carprofen was administrated s.c. post 24-hour surgery. Animal welfare was monitored by physical appearance, behaviour, and body weight. Power analyses were carried out prior to experiments being carried out to determine the minimum number of animals required for each experiment. These analyses were informed by previous and / or preliminary experiments. Animals were house vested in groups of four. Every group (i.e. cage) contained mixture of mice transplanted with both organoid genotypes. No animals were excluded from downstream analysis. The biotechnician was blinded to the group allocation of the animals during the experiment.

### Immunohistochemistry (IHC) of NSG mice tissues

Mouse tissues with or without organoid-initiated tumours/metastases were fixed in 4% formaldehyde solution and paraffin embedded. Sections of 4 µm thickness were made. Prior to the staining, tissue sections were deparaffinized and rehydrated. Endogenous peroxidase activity was blocked with 1.5% hydrogen peroxide for 30 min. Heat-induced antigen retrieval was carried out using citrate buffer pH6.0 for 20 min, followed by the cooling of tissue sections for 20 min. Sections were incubated overnight at 4 °C with primary antibodies (anti-hNucleoli clone NM95, Abcam (ab190710) 1:1000 and anti-Mucin2 clone (H-300), Santa Cruz (sc-15334) 1:100) diluted in PBS with 0.1% Sodium azide and 3% BSA. The next day, sections were washed three times with 0.05% Tween-PBS solution for 5 min, followed by a 1-hour incubation with the HRP-conjugated secondary antibody. Subsequently, sections were washed three times with 1× PBS for 5 min and developed with 3,3’-Diaminobenzidine (DAB) chromogen for 10 min at room temperature in the dark. Sections were rinsed under running tap water for 10 min and counterstained with hematoxylin, followed by dehydration and mounting.

Stainings were scanned using NanoZoomerXR whole slide scanner (Hamamatsu) at 40X magnification, with a resolution of 0.25 μm/pixel. Quantification of the scans was performed using QuPath program analysis program^61^. Target of interest (tumour or distinct immune populations) was quantified using QuPath’s Trained Pixel classification command that allows an automated recognition of background, tissue (hematoxylin) and DAB staining areas. The percentage of target of interest area was then calculated by using the following formula: DAB area / (DAB + tissue– background area) * 100.

### Reconstitution assays using growth factor and drug treatments

Single cells from organoid were harvested using TrypLE incubation at 37 °C. Cells were filtered through a 40 µm nylon cell strainer (Falcon). Cells were counted and seeded in a 1*10^6^ cells / mL dilution in Matrigel/BME. Cultures were incubated in standard culture medium with indicated additions/depletions of factors and drugs, which was replenished every 2-3 days. After seven days, brightfield images were taken using an EVOS imaging system and organoids were split 1:2 similar as mentioned before, after which outgrowth was monitored again for seven days. Quantitative viability assays were performed by CellTiter-Glo® (Promega) or Alamar Blue® (Thermo Fisher) according to manufacturer instructions. Pictures are shown for a representative experiment, which was performed at least three times. Normalised viability data was combined from multiple experiments and two or three technical replicates per experiment.

### Bulk RNA sequencing

Organoids were lysed in RLT lysis buffer (Qiagen) + 1% β-mercapto-ethanol; seven days after passaging. Biological triplicates were employed that were passaged and lysed at separate moments. For the LGK-974 experiments, organoids were treated with 100 nM LGK-974 or a DMSO volume control for 24 or 48 hours prior to lysis. RNA was obtained using the Qiagen QiaSymphony SP (Qiagen) according to the manufacturer’s protocol. Library prep was performed using Truseq RNA stranded polyA (Illumina). Libraries were sequenced on an Illumina NextSeq2000 platform with 1×50bp reads. Quality control on the sequence reads from the raw FASTQ files was done with FastQC (v0.11.8). TrimGalore (v0.6.5) as used to trim reads based on quality and adapter presence after which FastQC was used to check the resulting quality. rRNA reads were filtered out using SortMeRNA (v4.3.3) after which the resulting reads were aligned to the human reference genome (GCA_000001405.15_GRCh38_no_alt_plus_hs38d1_analysis_set) using the STAR^62^ (v2.7.3a) aligner. Followup QC on the mapped (bam) files was done using Sambamba (v0.7.0), RSeQC (v3.0.1) and PreSeq (v2.0.3). Read counts were then generated using the Subread FeatureCounts module (v2.0.0) with the GRCh38.106 gtf file as annotation. CPM and RPKM Normalised versions of the counts table were generated using the R-packages edgeR (v3.28) as well as a normalised version using DEseq2^63^ (v1.28). In each comparison, genes were selected with an absolute fold-change > 1.5 and p adjusted < 0.05. Generation of heatmaps and scatterplots was done in using the pHeatmap and ggplot2 R packages.

### GSEA and gene signatures

Gene set enrichment analyses (GSEA) were performed using the Broad Institute GSEA tool (software.broadinstitute.org/gsea/index.jsp) with default settings, including permutation by gene sets. HALLMARK gene sets were derived from the Molecular Signatures Database.^34^ Gene set for REG4^+^ deep crypt secretory cells (DCSs) was defined as human orthologues (identified by bioDBnet) of the >2 fold upregulated genes with a p-value < 0.05 in Reg^+^ cells compared to Lgr5^+^ cells that were previously reported.^23^ Gene sets for human colon specific cell types were derived from a published single-cell transcriptome study (Wang et al. table S2).^24^ Gene sets for HUB040 single cell clusters were defined by marker gene analysis using the ‘FindAllMarkers’ command in Seurat comparing one cluster to all other clusters using a wilcox test. Threshold for adjusted p-value was set at p < 0.05. The WNT signature was reported previously by Van der Flier *et al.*^64^

For the analysis of secretory genes, a list of secreted proteins was downloaded from human protein atlas (“Secreted proteins predicted by MDSEC”)^65^. Downregulated secretory genes were defined as: significantly downregulated in at least three comparisons and not upregulated in the other comparison (RC1 vs HUB040, RC2 vs HUB040, RC1 vs NC1, RC2 vs NC1)

### Single cell RNA sequencing and analysis of organoids

Single cells from organoids were harvested using TrypLE incubation at 37 °C, seven days after passaging. Cells were filtered through a 40 µm nylon cell strainer (Falcon). scRNA-seq was performed according to the Sort-seq protocol^66^. Libraries were sequenced on an Illumina NextSeq500 at paired-end 60– and 26-bp read length and 75,000 reads per cell.

Reads were mapped to the hg38 reference genome. Analysis was performed with the R-package Seurat^67^ (4.0.4). Cells with 400-50000 unique features and 1000-7000 total transcript counts were included in the analysis. Scores for gene signature expression were calculated using the AddModuleScore function.

Signature enrichment in marker gene lists was determined using the Enrichr tool^68^ (https://maayanlab.cloud/Enrichr/).

To analyse fraction of TINCs in *RNF43*-corrected and non-corrected clones, cells were filtered according to previously mentioned settings and for each cell a module score for the original TINC signature was calculated. Cells with a module score > 0.61 were scored as TINC, which corresponded to 16% of cells in the HUB040 naïve sample (which was the fraction of cells mapping to cluster 1 in the original analysis). Data was plotted using ggplot2.

To validate the TINC signature in a human cohort, published RNAseq data of tumour epithelial cells of colorectal cancer patients were used^10^. Twenty *RNF43* wildtype patients from iCMS2 cohort based on single-cell RNA sequencing and inferred CNV result and four *RNF43* mutant patients were selected. After selecting patients, supervised clustering using 154 TINC signature genes was performed with batch correction using harmony algorithm^69^ to remove patient specificity. Differential gene expression analysis was performed using the ‘FindAllMarkers’ command in Seurat with logistic regression framework.

### TCGA data analysis

Signature scores for indicated gene sets were calculated for each sample as log2 Z-score over the total TCGA Colon Adenocarcinoma set (rpkm values of 382 samples – cbcoad) using the R2 genomic analysis platform (http://r2.amc.nl). TINC signature was determined as the top 25 marker genes for cluster 1 in HUB040 (from Seurat marker gene analysis). *RNF43* mutation status was determined by combining the list of *RNF43* mutant TCGA samples (derived from C-bioportal) combined with results of re-analysis by Gianakkis *et al*.^22^ calling additional mutations that were falsely labelled as polymerase slipping errors in the original TCGA analysis.^22^ CMS classifications were derived from the Colorectal Cancer Subtyping Consortium (CRCSC)^40^, samples without CMS label have been left out of the analysis. Classification as mucinous was derived from Cancer Type in cbioportal (mucinous adenocarcinoma of colon and rectum).

### Immunofluorescence

For staining of Mucin2, organoids were seeded in 15-well IBIDI angiogenesis chambers during passaging and fixed 7-12 days later in 4% paraformaldehyde (PFA) (Sigma-Aldrich) for 60 min at room temperature. The fixation was quenched using 0.05 M NH4Cl and cells were permeabilised with PBD0.2T (48,5mL PBS, 1mL 10%Triton, 0,5mL DMSO and 0,5g BSA) for 20 min at room temperature. The organoids were incubated with primary antibodies diluted in PBD0.2T at 4 °C overnight. Washes were performed with PBD0.2T after which secondary antibodies were incubated for 3h at room temperature. Stained organoids were mounted using IBIDI mounting medium.

For other stainings, organoids were cultured as usual. 7-12 days after passaging, matrix was removed using dispase (Invitrogen) and organoids were transferred to 8-well IBIDI imaging slide. Fixation, permeabilisation and antibody staining were performed inside the imaging slide, as indicated above, except for primary antibody incubations were performed for one hour at room temperature. Stained organoids were mounted using IBIDI mounting medium.

The following antibodies and dyes have been used for immunofluorescence: Mouse-anti-Mucin2 (sc-515032, Santa Cruz, 1:200), Mouse-anti-β-catenin (610154, BD Transduction Laboratories, 1:200), Rabbit-anti-KI67 (ab15580, Abcam, 1:200), Goat-anti-Mouse-Alexa-488 (A11029, Invitrogen, 1:600) Goat-anti-Rabbit-Alexa-647 (A21245, Invitrogen, 1:600), Phalloidin-Alexa-647 (8940S, CST, 1:200) and Dapi (D9542, Sigma, 1:1.000). Images were acquired with a Zeiss LSM700 confocal microscope.

Image processing was performed using FIJI (2.3.0). Nuclear β-catenin and TINC-reporter intensity were quantified using the StarDist plugin (0.8.3)^70^ with the Dapi staining used for single nucleus segmentation. Data was analysed and plotted in R.

### Hybrid organoid co-culture, time-lapse imaging and analysis

The protocol on generating hybrid organoid co-cultures has been described previously^25^. In short: HUB040 naïve organoids were transduced with mCherry tagged H2B constructs and HUB040 RC2 organoids with mNeon green tagged H2B constructs and selected by FACS sorting. Six days post splitting, both lines were starved in CRC medium for 24 hours and at day 7 broken up into small clumps of cells that were mixed in 1:1 ratio. Cells were concentrated by centrifugation, resuspended in a small volume of CRC medium and incubated at 37 °C for 30 min Cell aggregates were plated in 10 μL BME droplets in an 18 Well µ-Slide (IBIDI #81811). Three days after plating, time-lapse microscopy was performed on an inverted LSM880 Fast AiryScan scanning confocal microscope (Zeiss) using LSM scanning mode with a 488nm Argon-laser and 561nm Diode-laser, equipped with a 20X dry immersion objective (Objective Plan-Apochromat 20x/0.8 M27, Zeiss). Cells were kept in CRC medium at 37°C under 5% CO2 for the full duration of the experiment. Images were acquired with a Z-interval of 5μm covering the full volume of the organoid and time-interval of one hour for a total duration on 60 hours. Time-lapse images were processed and analysed following a previously described pipeline in which depth-encoded movies were generated in which cell divisions and cell dead events were scored^71^. The cell death / mitosis ratio was calculated as a proxy for organoid survival. Cell numbers at the start of time-lapse were counted using Imaris software (Andor version 9.9.1). Videos and snapshots of 3D-reconstructed representative organoids were made in Imaris.

### Flow cytometry analysis and sorting of organoids

Single cells from organoid were harvested using TrypLE incubation at 37 °C. Cells were filtered through a 40 µm nylon cell strainer (Falcon) and live cells were measured at a FACS Celesta with FACS Diva software (BD). The following fluorescent channels were used: BB515 for mNeon, PE-CF594 for mRuby and PE-Cy5 for Alexa-647 labelled Edu.

EdU incorporation assays were performed by culturing organoids in CRC medium containing 10 µM EdU for 16 hours to cells, after which single cells were harvested, fixed in 4% PFA and EdU-labelling was performed according to manufacturer instructions (Click-iT EdU Cell Proliferation Kit for Imaging, Thermo Fisher).

Sorting of TINC-reporter populations was performed using a FACS Aria III machine (BD) Populations were gated for viable single cells which were divided in three equal populations of 20-30% of the total based on the mNeon distribution (example in Fig. 4e). Sorted cells were directly lysed in RLT lysis buffer (Qiagen) for RNAseq analysis, or taken in culture for organoid reconstitution, by seeding in a 1×10^6^ cells / mL dilution in BME.

Gated single cells were analysed in FlowJo (version 10.6.1).

### Western blotting

Organoids were collected in Cell Recovery Solution (Corning), incubated for 20 min on ice and subsequently lysed in RIPA buffer containing protease and phosphatase inhibitors. Lysates were boiled in 1x SDS sample buffer and ran on a pre-casted SDS-page gel (Mini protean, BIO-RAD). Gels were transferred to a Immobilon-FL PVDF membranes (Millipore), which was blocked for 1 hour at room temperature in 1:1 ratio Odyssey blocking buffer (LI-COR): PBS, and stained against APC (sc-7930, Santa Cruz, 1:1.000) and Actin (691001, MP, 1:10.000) overnight at 4 °C. Blots were stained with secondary antibodies IRDye-680 goat anti-rabbit (926-32221, LI-COR, 1:10.000) and IRDye-800 goat anti-mouse (925-68070, LI-COR,

1:10.000) for 1 hour at room temperature in the dark. The Typhoon (GE Healthcare) infrared imaging system was used for immunoblot analysis.

### RT-qPCR

Organoids, seven days after passaging, or FACS-sorted cell populations were lysed in RLT lysis buffer (Qiagen) + 1% β-mercapto-ethanol and RNA was purified using the RNeasy kit (Qiagen) according to manufacturer’s protocol. RNA was used as a template for cDNA production using the iScript™ cDNA Synthesis Kit (Biorad) according to the manufacturer’s protocol. The synthesized cDNA was subsequently used in a qRT–PCR using IQ SYBR green mix (Biorad) according to the manufacturer’s protocol. Used primers are listed in Supplementary Table 2. Expression of WNT target genes was corrected against house-keeping gene *GAPDH* and normalised to the control.

### iCaspase experiments

Imaging: Organoids were seeded in an 8-well Labtek imaging chamber and after 6 days organoids were imaged overnight (15 min interval) using the Nikon Spinning Disk microscope. Right before imaging DMSO or 10 nM AP20187 (Sigma Aldrich SML2838) was added to the organoids. Movies were annotated using a plugin in Fiji^72^.

For the cell viability assay, single cells were seeded into 8-wells of a 48-well plate (∼2000 cells/well) and grown in CRC or HCO medium supplemented with either DMSO or 10 nM AP20187 for 14 days. Medium was refreshed every 2-3 days and images were taken on an EVOS microscope. At day 14 cell viability was assessed by a CellTiter-Glo assay (Promega) according to manufacturer’s protocol.

### APC competition assay

HUB040 and HUB040^APC-/-^ organoids were split into single cells, counted and seeded in a 1:1 ratio. Mixed organoid populations were passaged once a week, and during each passage, genomic DNA was harvested for genotyping the fraction of *APC* mutant alleles as indicated above.

### Mouse model studies

Mouse studies were carried out in accordance with the UK Home Office Guidelines (project licence 70/9112) under approval of the University of Glasgow Animal Welfare and Ethical Review Board. Mice were housed in a pathogen-free facility in conventional open-top cages in compliance with UK Home Office Regulations, straw bedding material and tunnels were available in all cages as enrichment. Mice were maintained at constant temperature and humidity under a 12-hour light/dark cycle with access to a standard chow diet and water ad-libitum. All mice were genotyped by ear punch samples obtained at time of weaning and processed externally (Transnetyx, Cordova, TN). At age of weaning, a maximum of 5 male or female mice were kept in the same cage. The use of single-housed experimental animals was avoided throughout this study. The following transgenes and alleles were used within this study: VillinCreER^73^, *BrafV600E*^74^, *Trp53fl/fl*^75^, *Rnf43fl/fl*^5^. Descriptions of the genetically engineered mouse models (GEMMs) developed in the CRUK Scotland Institute (Sansom Lab) are detailed in Supplementary Table 3.

Experimental mice were maintained on a C57BL/6 background (n ≥ 5). Experimental mice received a single 2 mg intraperitoneal injection of Tamoxifen (Sigma-Aldrich T5648). Mice were induced between 6 and 14 weeks of age at a weight of > 20 g, both male and female mice were used. Mice were weighed at a minimum of 3 times per week and monitored daily for signs of intestinal tumourigenesis. Humane clinical endpoints were determined based on the presence of weight loss and loss of body condition and/or abdominal swelling and/or hunching and/or presence of paling feet. Mice that were identified with health reasons not related to intestinal tumour burden (including lymphoma and skin wounds) were censored.

### Tissue sampling, processing and staining of mouse models

On clinical endpoint, mice were culled and samples collected in 10% neutral buffered formalin for 24-72 hours. Whole small intestine (SI), colon and caecum were removed and flushed with water prior to opening the gut longitudinally and pinning on a wax plate with the intestinal lumen facing upward. The mesenteric lymph-nodes, liver, lung, pancreas and spleen in addition to any other identifiable organs with metastatic disease at time of dissection were also collected. Following formalin fixation, intestinal tissues were rolled into a “Swiss roll”. Both the intestinal tissue and organ samples were transferred to 70% ethanol with organs.

Tissue samples were processed and embedded for Haematoxylin and Eosin (H&E) staining by the CRUK Scotland Institute Histopathology Department under standardised techniques. The process of sample collection and histological scoring of samples was performed in a blinded manner. For Muc5a staining, mouse anti-Muc5ac (ab3649 Abcam, 1:100) was used. Slides were scanned and digitalised using the Lecia SCN400F slide scanner at a 20X magnification. To generate representative histological images of tumours HALO software v2.0 (Indica Labs) was used.

### Single cell RNA Sequencing mouse models

A total of 40,000 cells were loaded onto each channel of Chromium Chip G using reagents from the 10x Chromium Single-Cell 3’ v3 Gel Bead Kit and Library (10x Genomics) according to the manufacturer’s protocol. The libraries were analysed using the Bioanalyzer High Sensitivity DNA Kit (Agilent Technologies). scRNA-Seq libraries were sequenced on the Illumina NovaSeq 6000 with paired-end 150-base reads. Sequence alignment of single cell data to the mm10 genome was performed using the count tool from the Cellranger package version 6.1.2 according to the developer’s instructions, generating barcodes, features and matrix output files for each sample. Subsequent analysis was done using R version 4.1.1 using Seurat version 4.0.4. Samples were input using the Read10X function, filtering to include cells with a minimum of 100 expressed genes and genes that are present in at least 3 cells, then further filtered to only include cells with <5% mitochondrial genes, <10% hemoglobin genes, >100 genes/cell and >400 reads/cell. Samples were then integrated by RPCA using the IntegrateData function before being scaled and normalised. Dimension reduction was then performed using PCA before clustering was performed using the FindNeighbours and FindClusters functions. Epithelial cells were isolated using CellTypist with the ‘Adult Mouse Gut’ template, and Epcam positivity, then reclustered. Gene expression module scores were added using the ‘AddModuleScore’ function.

## Statistical analyses

For RNA sequencing analysis of organoids, statistical significance was determined using the DESeq function in R, which uses a Wald test (two-tailed), see detailed methods. Gene set enrichment was performed as described in the methods. For other data, p values were calculated using the ‘stat_compare_means’ function of the ggpubr package (0.40.0) in R. For comparing two or more groups of non-parametric data, a two-sample Wilcoxon test or Kruskal-Wallis test were used respectively. For mouse model studies, a priori sample size calculation was performed based on historical data sets to ensure that the smallest experimental sample size deployed within the study could yield a significant difference in accordance with the 3Rs and accounted for the occurrence of any censored animals within the study.

Statistical analyses were performed using GraphPad Prism V10 for Windows. Survival analysis of BP BRNF and BPRNF models was performed using Kaplan-Meier plots and compared for significance using the log-rank (Mantel-Cox) test. p ≤ 0.05 was considered significant.

## Data availability

The scRNA-seq and bulk RNA-seq data of organoids generated in this study are available from Gene Expression Omnibus (GEO) under the accession code GSE226256. BP and BPRNF single cell RNAseq data is available upon request and will be deposited before publication.

Published scRNA-seq data referenced in the study is available from Synapse under the accession code syn26844071. Published microarray data are available from Gene Expression Omnibus (GEO) under the accession code GSE41258.

## Acknowledgements

The authors thank members of the Maurice laboratory for helpful discussions and suggestions. We thank Bas Ponsioen for help with time-lapse image analysis. This work is part of the Oncode Institute, which is partly financed by the Dutch Cancer Society. This work was supported by the Netherlands Organization for Scientific Research NWO VICI Grant 91815604 (to M.M.M) and ZonMW TOP Grant 91218050 (to M.M.M., O.K. and H.J.G.S.), the IMAGINE! Gravitation grant 024.005.009 (to M.M.M. and H.J.G.S.), Dutch Cancer Society Young Investigator Grant 11491/2018-1 (to S.J.E.S.), NWO Rubicon Grant 019.233EN.017 (to J.M.B.), the Cancer Research UK core funding to the CRUK Scotland Institute (to O.S) and a CRUK Senior Group Leader programme award (A29055, A28223, A21139, DRCQQR-May21\100002, CTRQQR-2021\100006]DRCQQR-May21\100002) (to O.S), a Cancer Research UK Accelerator Award (A26825) and a Mark Foundation ASPIRE I Award (to M.L.M., A.C., and O.S.).

## Author Contributions

J.M.B., L.E.B., E.K., H.J.G.S., O.K. and M.M.M. conceptualized the project. J.M.B., O.S., O.K. and M.M.M. designed and interpreted experimental results. J.M.B., L.E.B., E.K., M.L.M., J.S., I.J., D.K., S.P., A.V., S.B., and S.J.S. performed experiments and analysed results. Y.H., K.G., D.M.G., R. V. B. and S. T. provided support with data analysis. J.H. and N.F. provided methodology. J.M.B. and M.M.M. wrote the manuscript, which was reviewed by all authors.

## Extended Data Figures

**Extended Data Figure 1.**
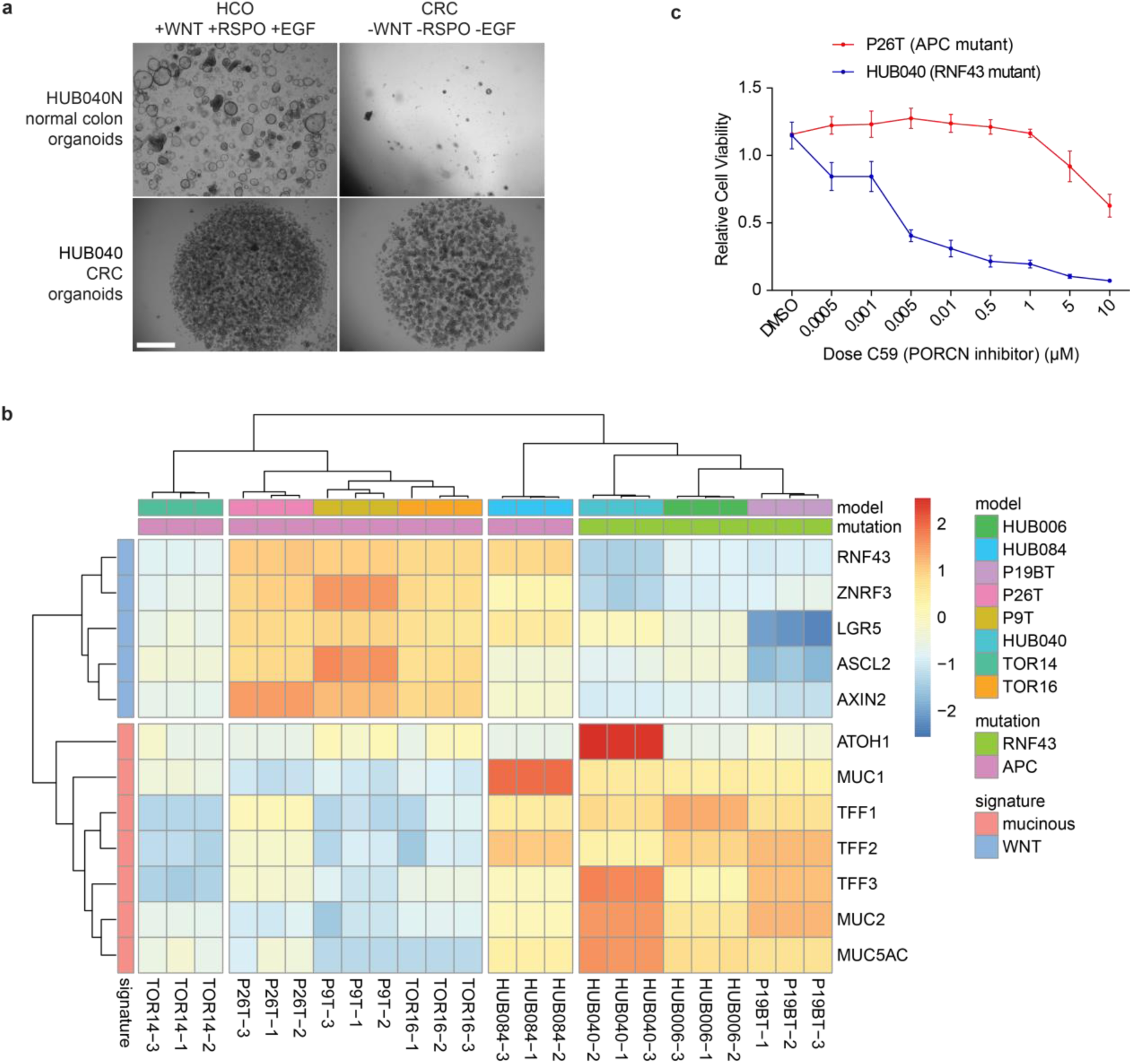
| Patient-derived *RNF43*-mutant CRC organoids are growth factor self-sufficient. **a.** Representative brightfield images showing healthy colon organoids and HUB040 CRC organoids cultured in CRC culture medium or complete colon organoid medium (including WNT, EGF and RSPO) for e passage and 14 days in total. Scale bar = 1000 μm. **b.** Heatmap of gene expression data for WNT markers or mucinous markers over a panel of CRC organoids (n = 8). Presence of *RNF43* or *APC* mutation is indicated on top. Unsupervised clustering into four groups shows that *RNF43*-mutant organoids cluster together in a low-WNT, high-mucinous group. Coloured bar represents row z-scores of normalised counts. **c.** Line graphs showing normalised viability of HUB040 and P26T organoids treated with indicated concentration of PORCN inhibitor C59 for 7 days. Data were normalised to PBS-treated organoids. n = 2, mean +/− S.D.

**Extended Data Figure 2.**
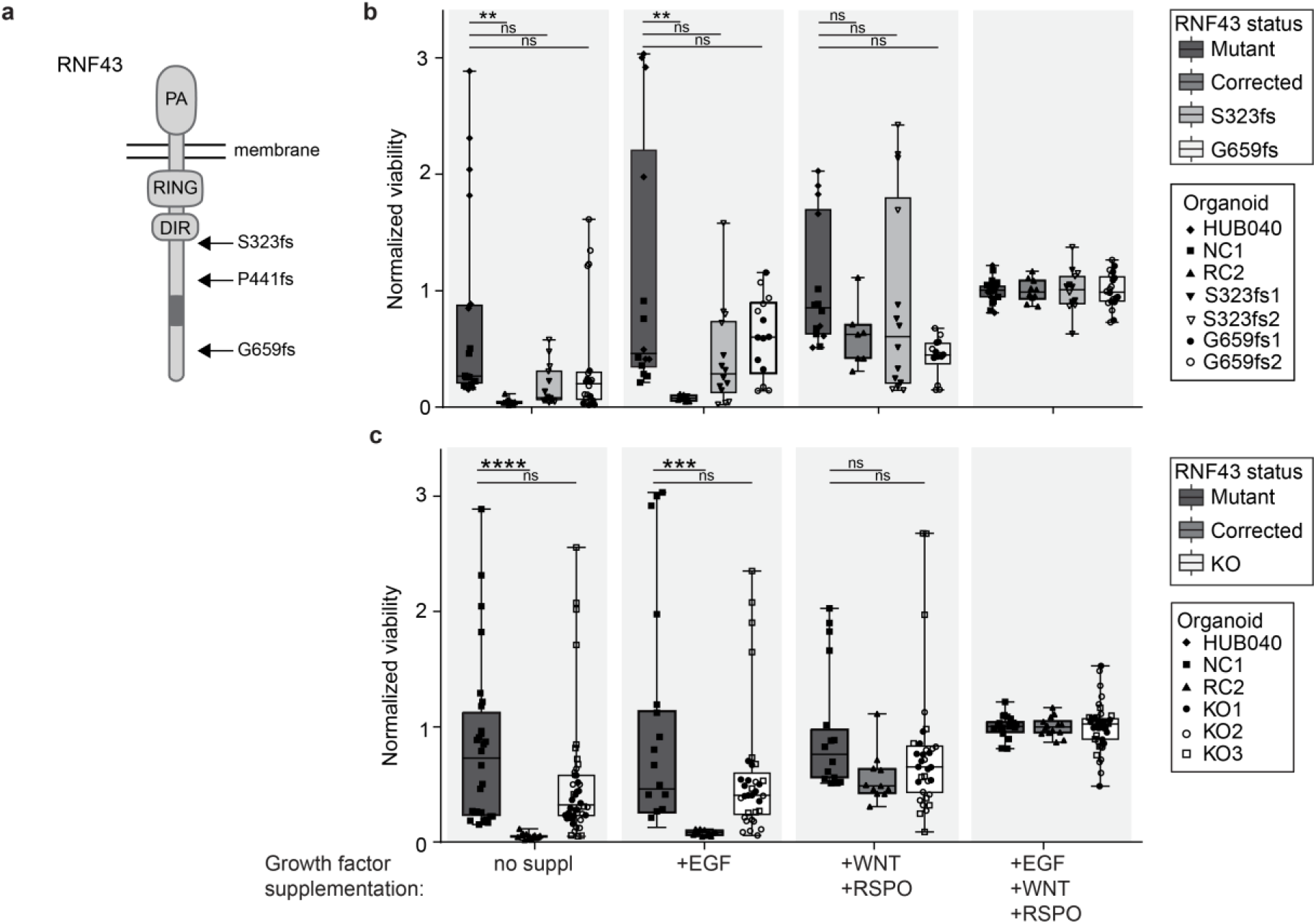
| Diverse subsets of RNF43 mutations facilitate niche factor independent growth. **a.** Schematic of the human RNF43 protein indicating the location of the S323fs, P441fs and G659fs mutations. PA: protease associated domain, RING: E3-ligase domain, DIR: Dishevelled-interacting region, dark grey: oncogenic region. **b, c.** Normalised viability of *RNF43* mutant, *RNF43*-corrected or either *RNF43*-corrected RNF43 S323fs or G659fs re-mutated (**b**) or RNF43 knockout (KO) (**c**) HUB040 clones cultured in CRC medium supplemented with indicated growth factors for 14 days, passaged after 7 days. Results of two to four experiments are presented, including three to four replicates per condition (organoid line in a given medium) per experiment (n = 7-14 per condition in total). Data are normalised against mean value in complete medium per experiment. Kruskal-Wallis test is used to compare groups to *RNF43* mutant within each condition. ns, not significant, ** (P < 0.01), *** (P < 0.001), and **** (P < 0.0001).

**Extended Data Figure 3.**
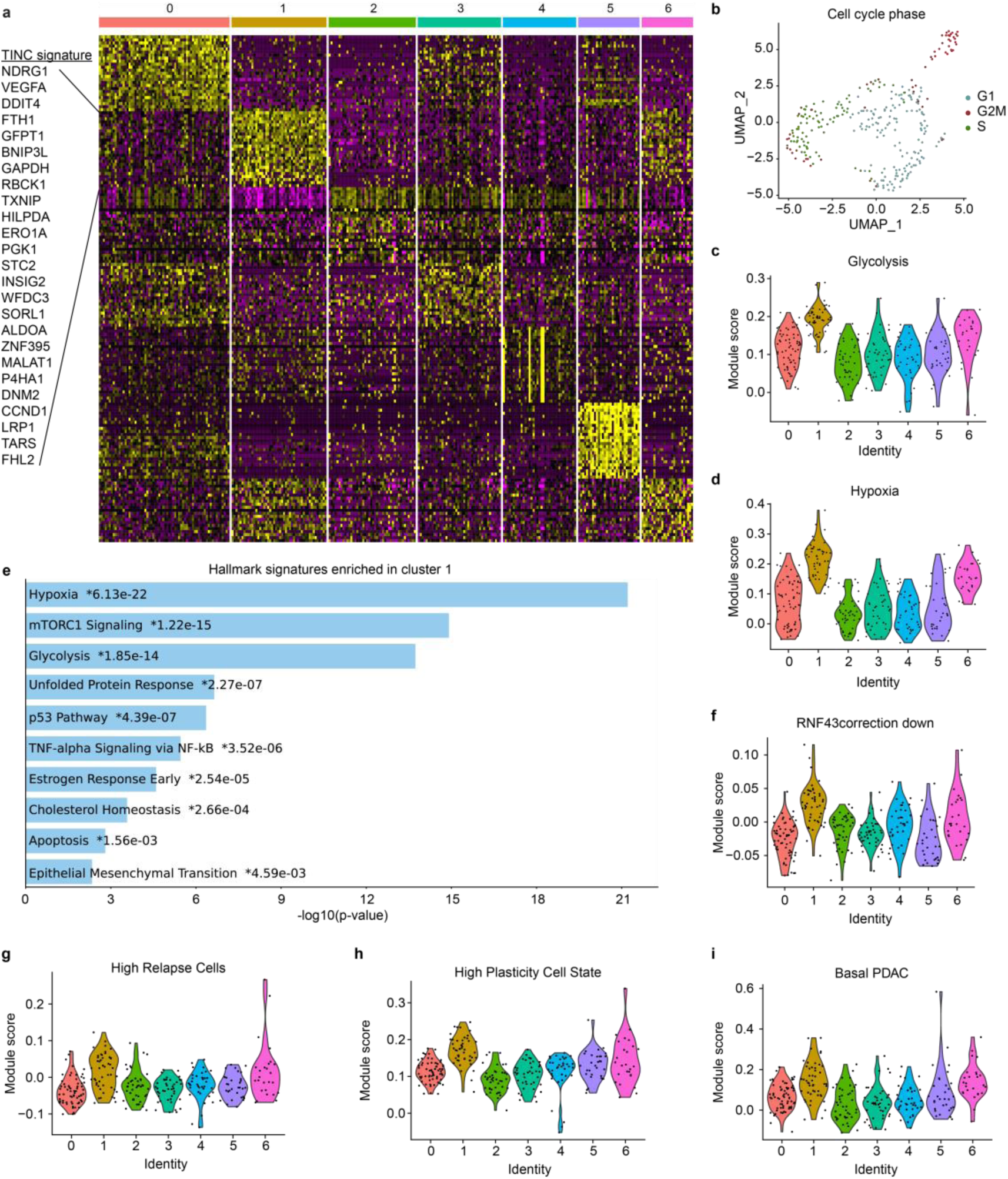
| Single cell RNAseq HUB040. **a.** Single cell expression heatmap showing row Z-scores for top 25 marker genes for each cluster. Marker genes for cluster 1 are highlighted and later used as ‘TINC signature’. **b.** Reduced dimensionality (UMAP) visualisation with cell cycle phase as determined by Seurat Analysis. **c, d.** Violin plots showing single cell module scores for expression of genes from the HALLMARK glycolysis (**c**) and hypoxia (**d**) signatures^34^ grouped by cluster. **e** Bar plot showing p-value of the top ten enriched HALLMARK gene signatures in the TINC marker genes list. **f.** Violin plots showing single cell module scores for expression of genes downregulated in RNF43 corrected clones grouped by cluster. **g-i.** Violin plots showing single cell module scores for expression of genes from the HRC^33^, HPCS^35^ and Basal PDAC signatures^34^ grouped by cluster.

**Extended Data Figure 4.**
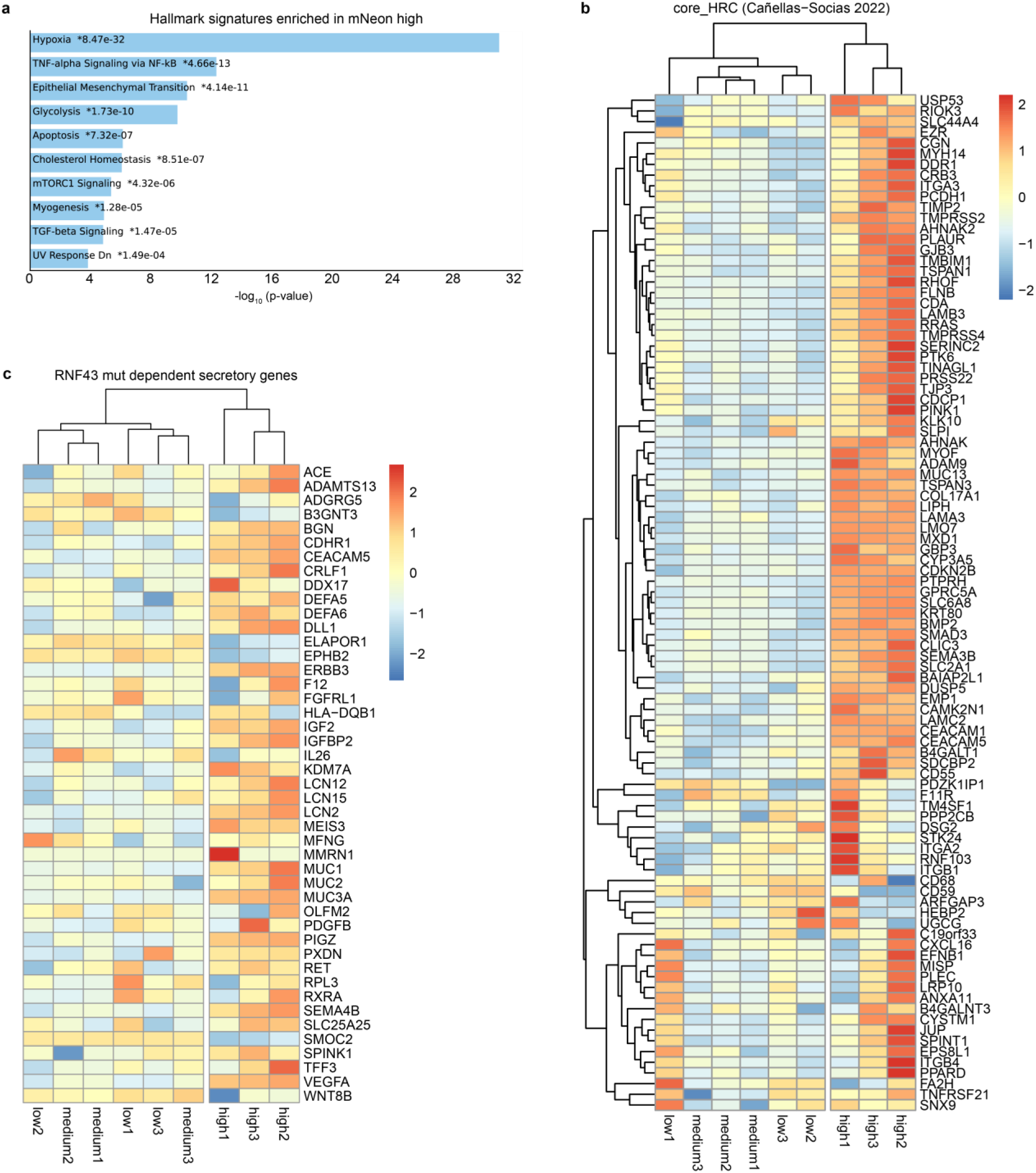
| Transcriptional characterisation of TINC reporter cells. **a.** Bar plot showing p-value of the top ten enriched HALLMARK gene signatures in the mNeon high group compared to the other groups. **b, c.** Heatmap of genes enriched in High Relapse Cells (HRC) (**b**) or downregulated secretory genes in *RNF43*-corrected HUB040 (as defined in Supplementary Fig. 3) (**c**) for the sorted TINC reporter populations. Coloured bar represents row z-scores of normalised gene counts.

**Extended Data Figure 5.**
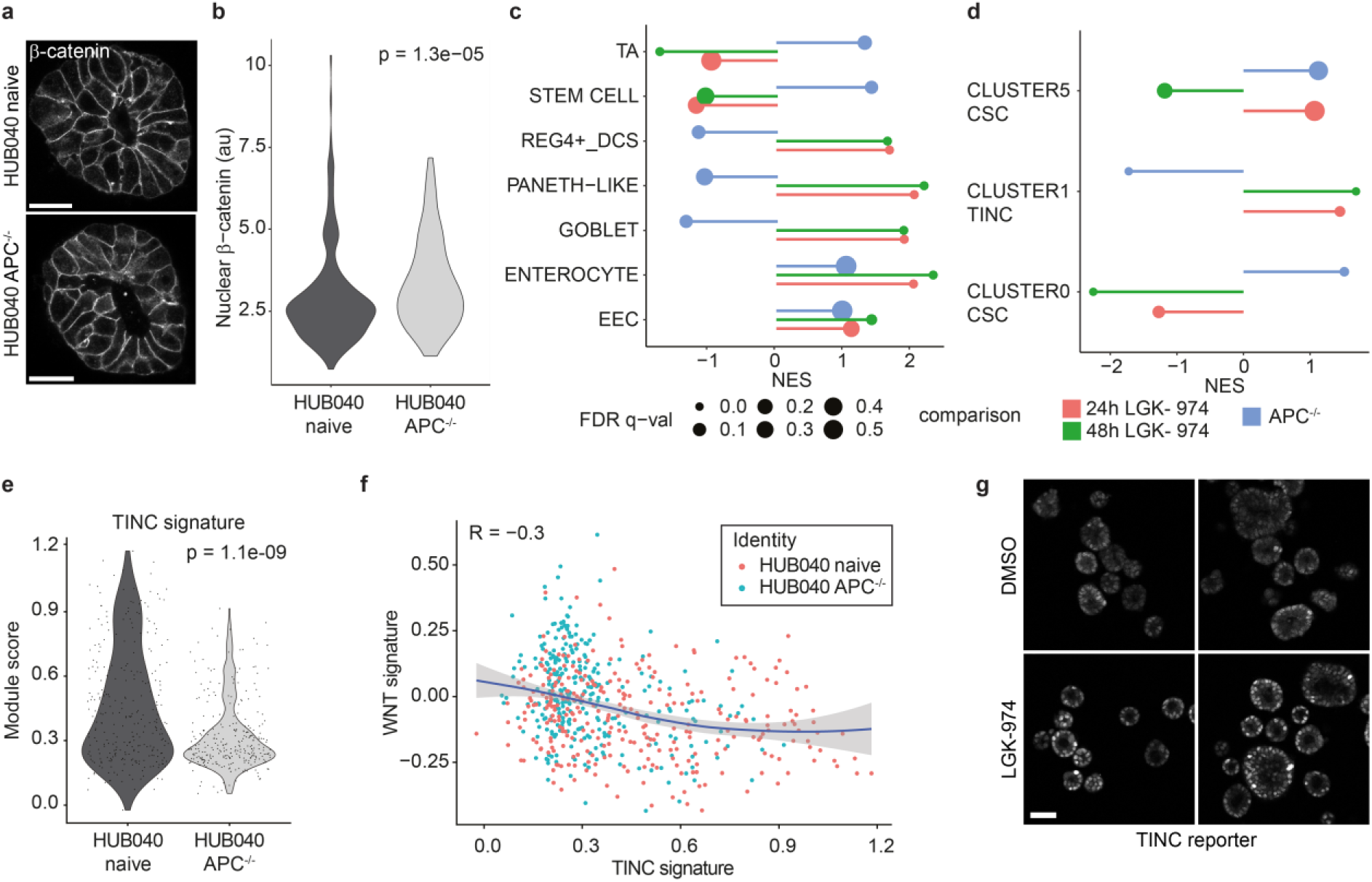
| A tight window of WNT signalling balances tumour niche cells and stem cells. **a.** Representative confocal microscopy pictures of organoids with IF labelling for β-catenin. Scale bar = 20 µm. **b.** Violin plot showing quantification of nuclear β-catenin signal for 205-231 cells per organoid line. **c, d.** GSEA analysis of HUB040 versus HUB040 *^APC-/-^* (WNT activation, blue), and HUB040 treated with DMSO versus HUB040 treated with 100 nM LGK-974 for 24 or 48 hours (WNT inhibition, pink and green) by gene signatures of colon-specific cell types^23,24^ (**c**) or HUB040-specific clusters as determined in Figure 3A (**d**). X-axis shows the normalised enrichment score (NES) and size of the dots shows the false discovery rate (FDR). **e.** Violin plots showing single cell module scores for expression of genes from the HUB040 TINC signature grouped by organoid line. **f.** Scatter plot showing inverse corelation between single cell module scores for expression of genes from the HUB040 TINC signature and the Van der Flier WNT signature^64^. Colour of dots indicates line of origin. **g.** Representative confocal microscopy images of HUB040 NDRG1-IRES-mNeon (TINC reporter) organoids after three days treatment with DMSO or LGK-974 (100mM). Scale bar = 50 µm.

**Extended Data Figure 6.**
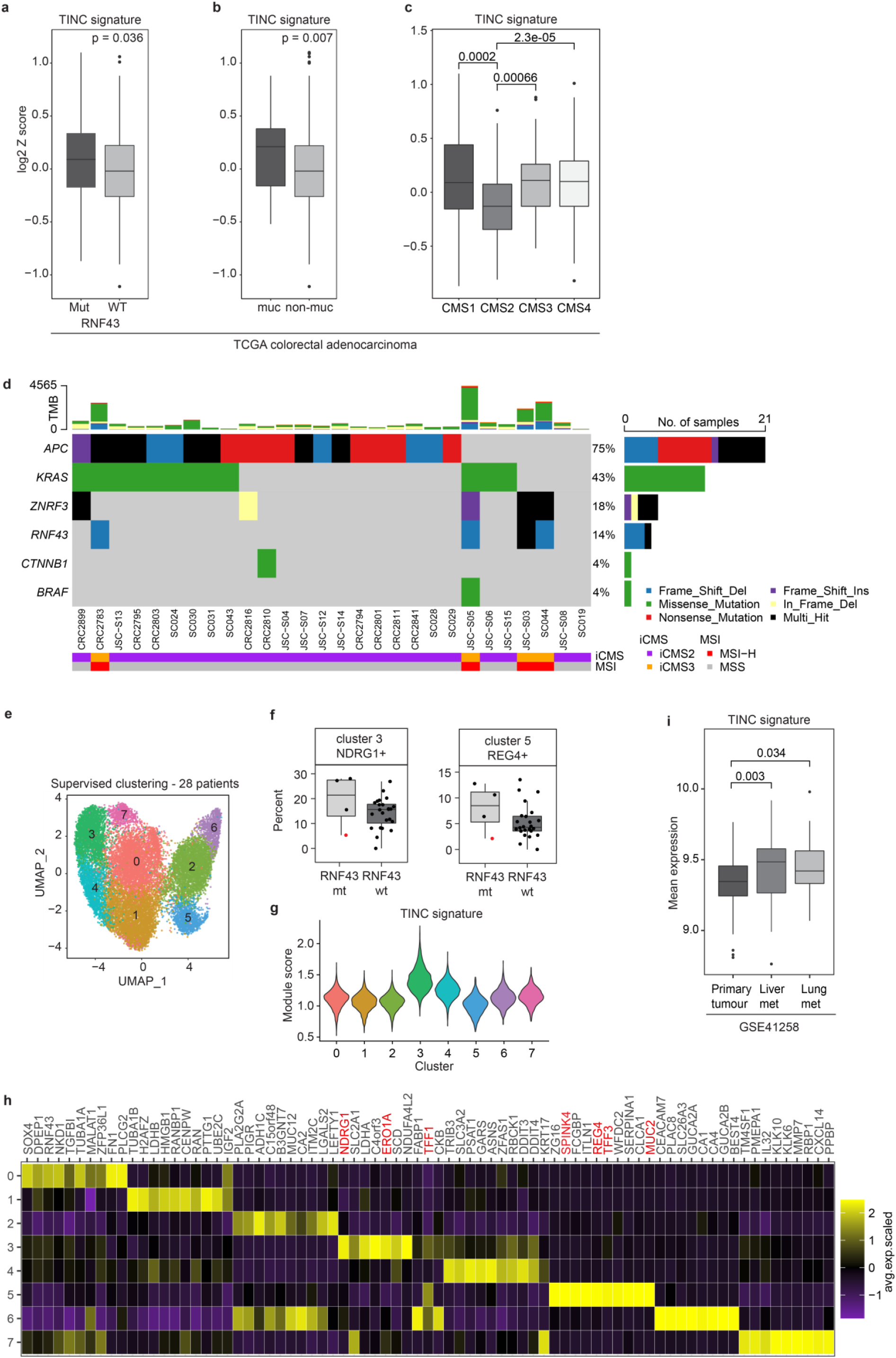
| TINCs are enriched in *RNF43* mutant, mucinous and metastatic CRC. **a-c.** Box plots showing log2 Z-scores for the TINC signature (top 25 differentially expressed marker genes of HUB040 cluster 1 as determined in single cell RNAseq analysis) over the colon adenocarcinoma TCGA bulk RNA sequencing dataset grouped by *RNF43*-mutation status (n = 46 Mut, 337 WT) (**a**), pathology classification as mucinous (n = 37) or non-mucinous (n = 346) (**b**) or consensus molecular subtype (CMS) as reported by Guinney et al^40^ (**c**). **d.** Oncoplot showing driver mutation status of the 28 patients included in single cell sequencing cohort^10^. **e.** Reduced dimensionality (UMAP) visualisation of supervised clustering using 154 TINC signature genes of 18.320 single cells from 28 patients by Joanito et al^10^. **f.** Box plots showing the fraction of cells in cluster 3 or 5, divided by *RNF43* mutation status per patient. The red dot marks a patient that has an *RNF43* and *APC* mutation. **g.** Violin plot showing single cell module scores for expression of genes from the TINC signature grouped by cluster. **h.** Marker gene analysis of the single cell patient data. Genes of interest are coloured red. **i.** Box plots showing expression of TINC signature (RMA normalisation data) in microarray data of 186 primary tumours, 48 liver metastases and 20 lung metastases^76^. Dots indicate outliers.

## Supplementary Information

**Supplementary Figure 1.**
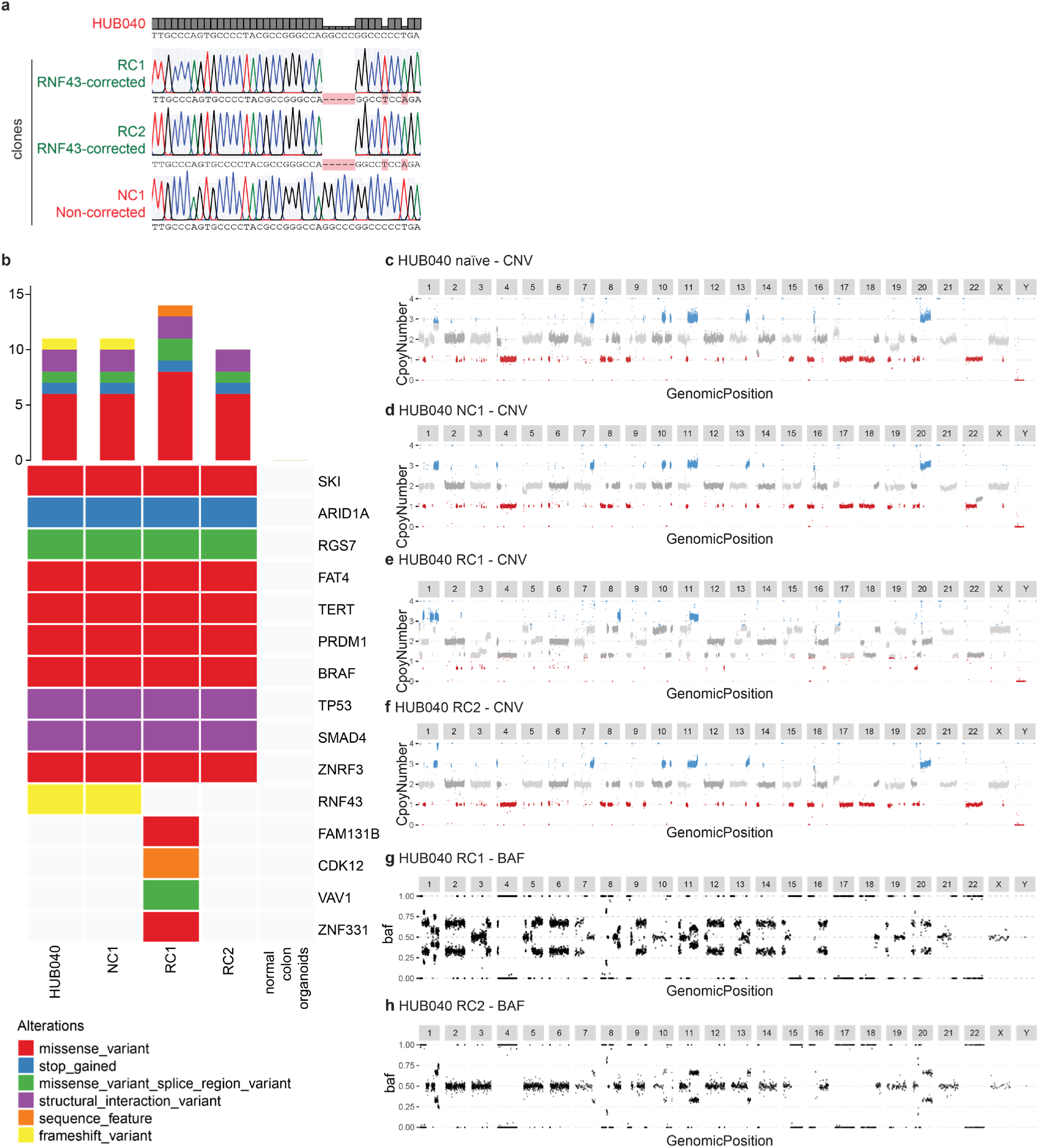
| Sanger and Whole genome sequencing analyses of HUB040 and clonal organoid lines. **a.** Sanger sequencing traces of RC1, RC2 and NC1 compared to the mutation locus of the HUB040 CRC organoid. **b.** Oncoplot showing high-confidence putative driver mutations derived from whole genome sequencing data of HUB040, normal colon organoids, NC1, RC1 and RC2. **c-f.** In silico karyograms of HUB040 naïve CRC organoids (**c**), Non-corrected control HUB040 clone (NC1) (**d**), *RNF43*-corrected clone 1 (RC1) (**e**) and *RNF43*-corrected clone 2 (RC2) (**f**). **g, h.** B-allele frequency (BAF) plot for RC1 (**g**) and RC2 (**h**) which is indicative of a whole genome duplication event for RC1.

**Supplementary Figure 2.**
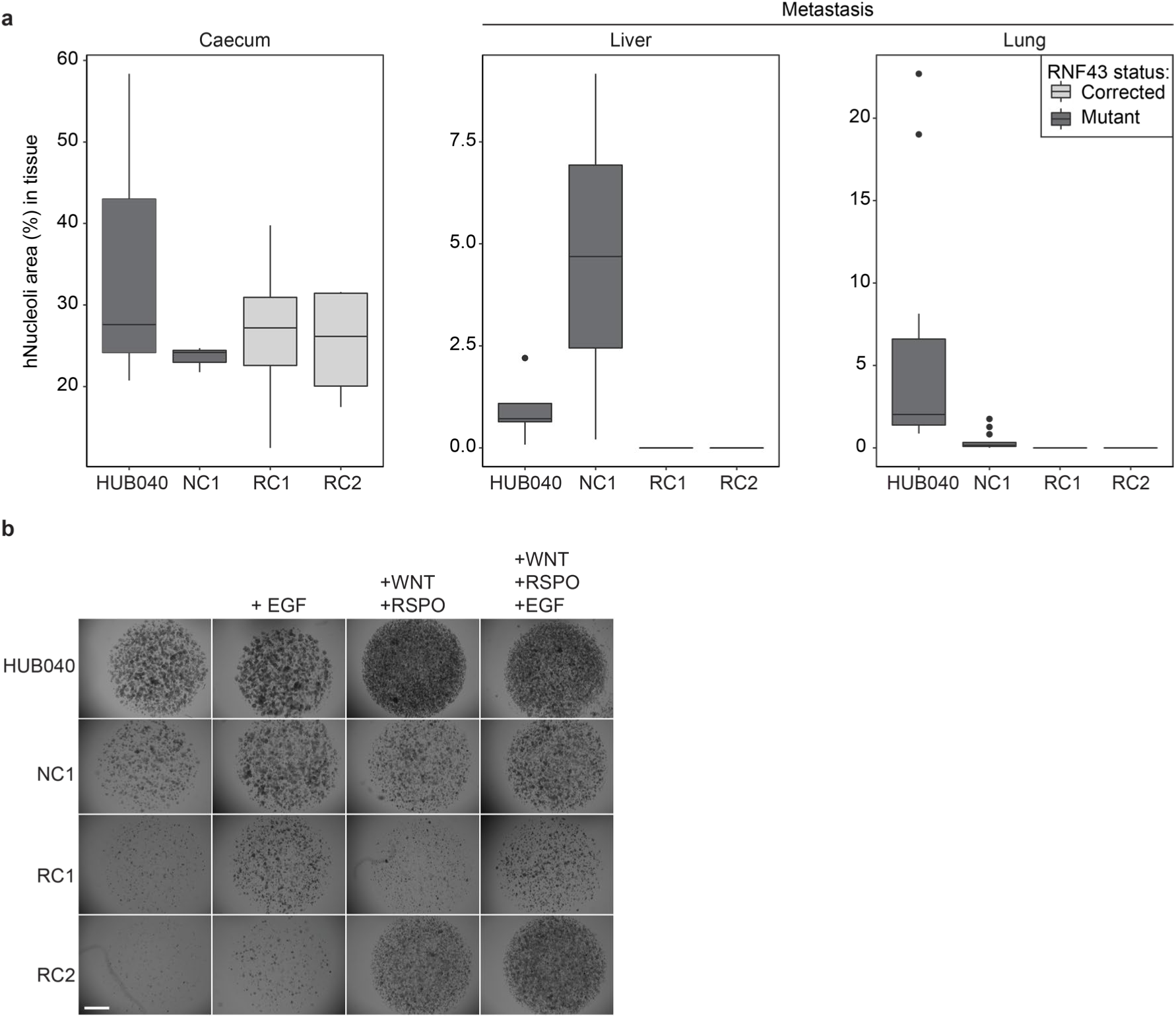
| RNF43 mutation facilitates niche factor self-sufficiency. **a.** Box plots showing the fraction of tumour tissue relative to total tissue area for caecum, liver and lungs in mice that had a tumour or metastasis (see Figure 1C for statistics). **b.** Representative low-magnification (2x) brightfield images of HUB040-derived clones cultured in CRC medium supplemented with indicated growth factors for 14 days, passaged after 7 days. Scale bar = 1000 μm.

**Supplementary Figure 3.**
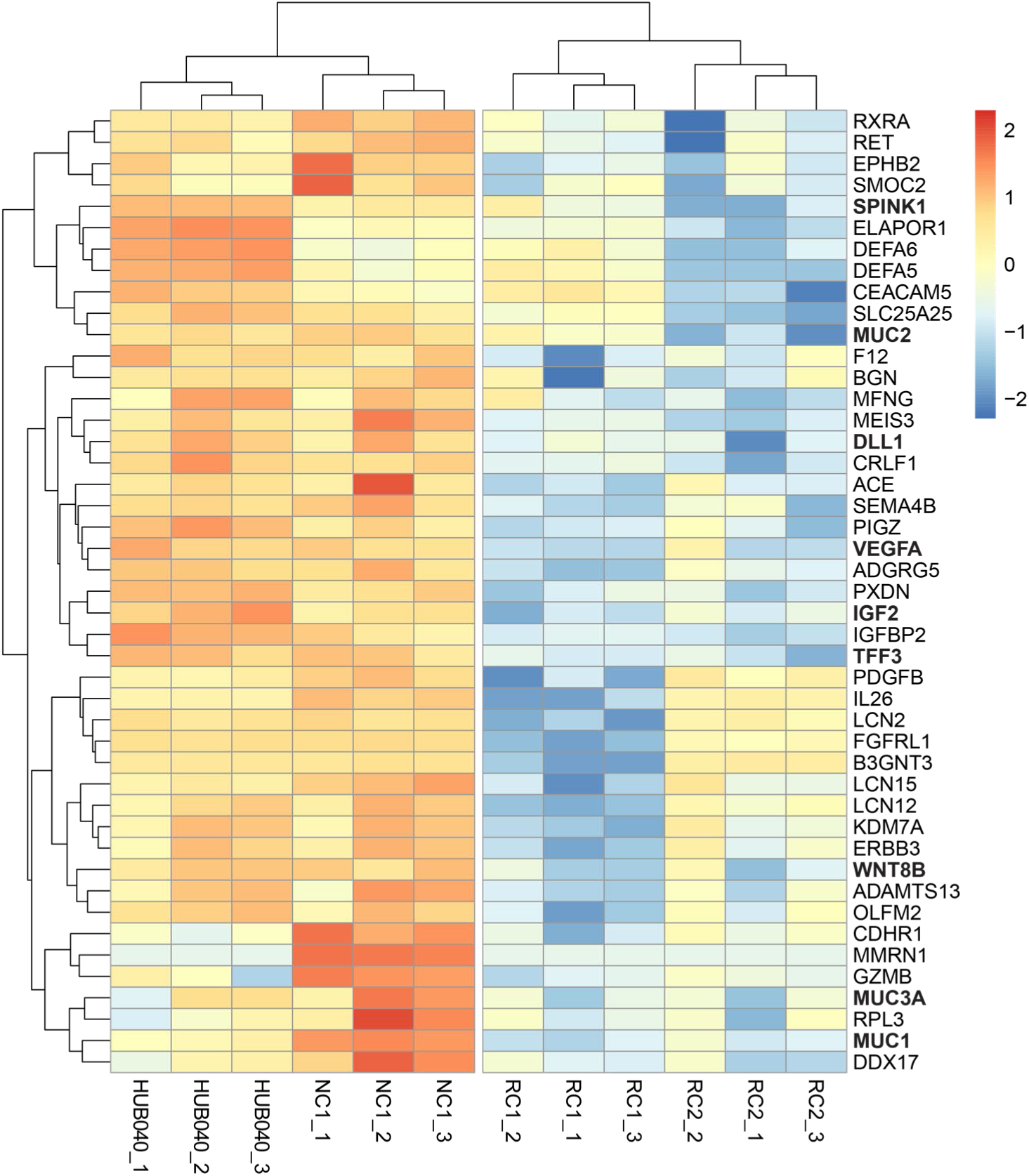
| RNF43 mutation upregulates secretory genes. Heatmap of downregulated secretory genes in *RNF43*-corrected HUB040 from a list of 3000+ genes encoding for secreted proteins, as predicted by MDSEC.^64^ Coloured bar represents row z-scores of normalised gene counts.

**Supplementary Figure 4.**
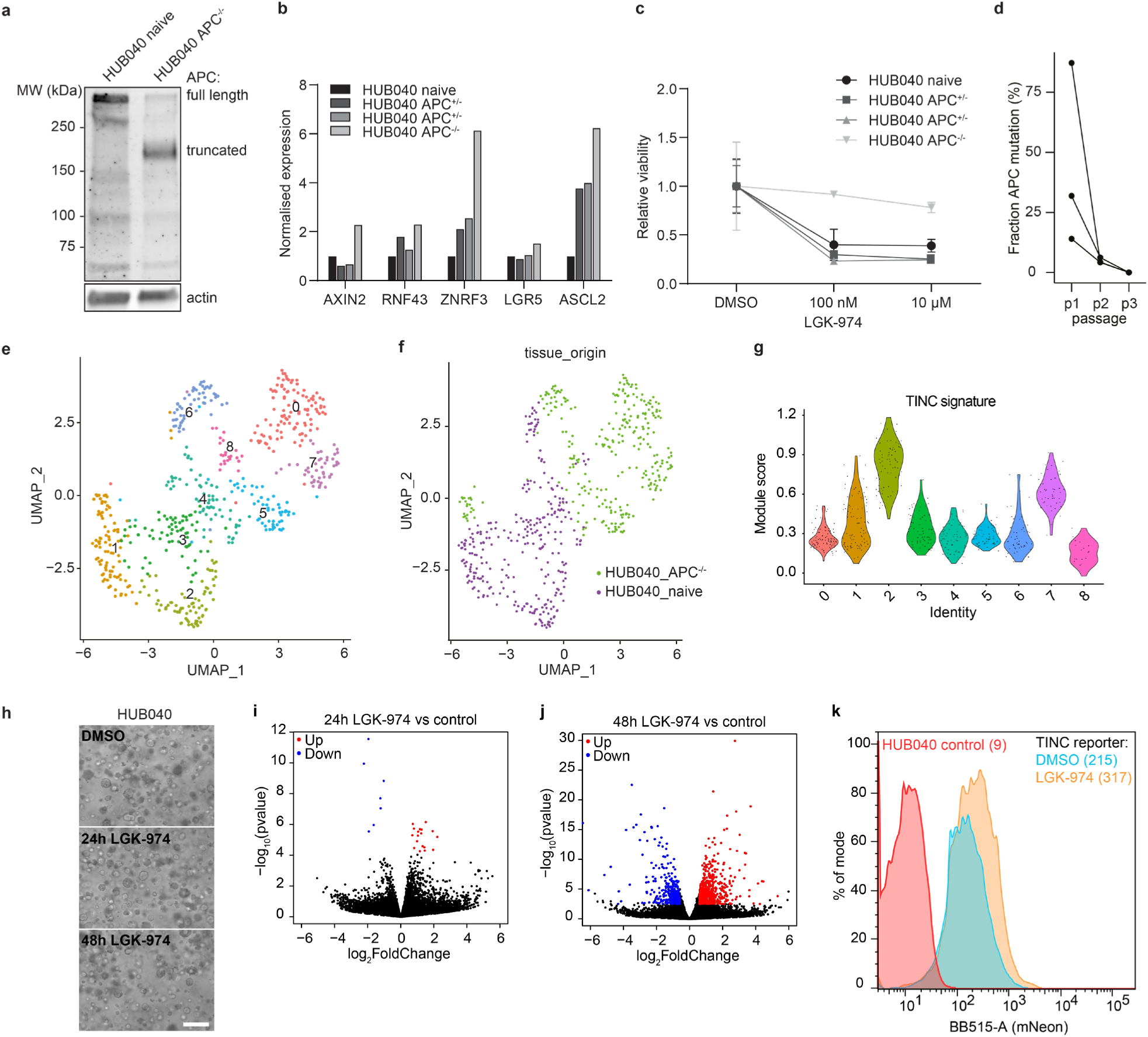
| Effects of APC mutation and LGK-974 on HUB040. **a.** Western blot showing truncated APC protein expression in homozygous *APC* mutant HUB040. **b.** Bar plot showing expression values, as measured by RT-qPCR, of WNT target genes in HUB040 organoids with CRISPR induced heterozygous (two clones) or homozygous (one clone) *APC* mutation normalised to naïve HUB040. **c.** Line graph showing relative viability of naïve HUB040 organoids, or HUB040 with heterozygous or homozygous *APC* mutation after seven days treatment with indicated concentration of PORCN-inhibitor LGK-974 or DMSO control. Dots indicate mean +/− S.D. n = 3. **d.** Line graph showing fraction of APC mutant DNA after each passage in the competition assay of indicated lines. n = 3. **e**. Dimension reduction (UMAP) plot of single cell RNA sequencing data of naïve and *APC* mutant HUB040 organoids. Results of unsupervised clustering are colour-coded per cluster. **f.** UMAP colour-coded per organoid line of origin. **g.** Violin plots showing single cell module scores for expression of genes from the HUB040 TINC signature grouped by cell cluster. **h.** Representative 10x brightfield images of organoid cultures used for RNA isolation. Organoids were treated with 100 μM LGK-974 for 24 or 48 hours, or with vehicle control (DMSO). Scale bar = 200 μm. **i, j.** Volcano plot of differential gene expression between 24h (**i**) or 48h (**j**) LGK-974 treated HUB040 organoid versus DMSO treated HUB040 organoids. Significantly (FC > 1.5 and p < 0.05) up and down regulated genes are plotted as red and blue dots respectively. **k.** Histogram showing FACS analysis of HUB040 NDRG1-IRES-mNeon reporter organoids after three days treatment with LGK-974 or DMSO control. Mean fluorescence intensity is indicated within brackets.

**Supplementary Table 1:**
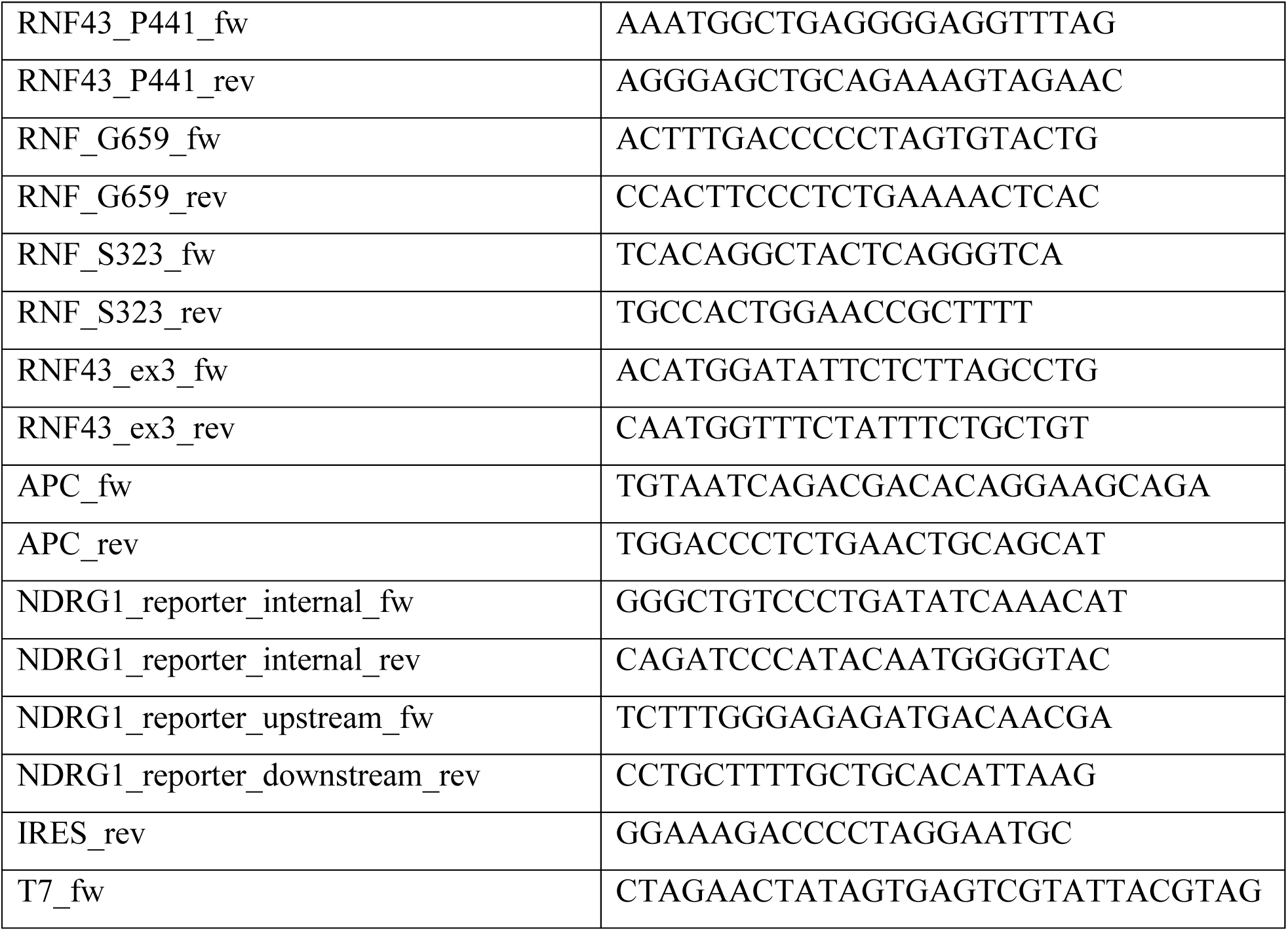
primers used for genotyping and Sanger sequencing.

**Supplementary Table 2:**
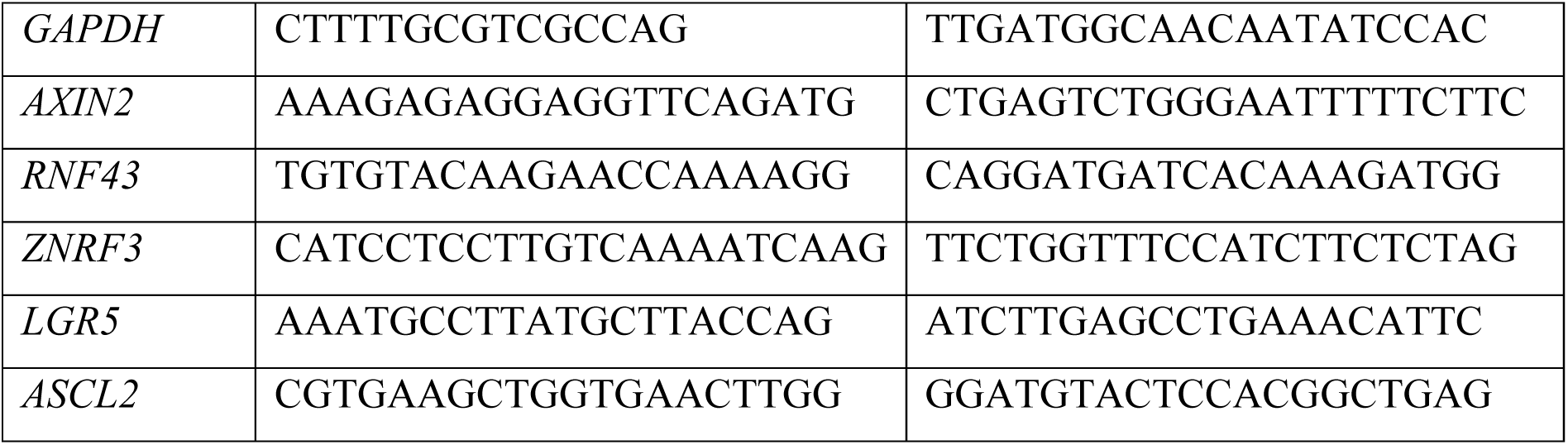
RT-qPCR primers.

**Supplementary Table 3:** Summary of GEMMs from CRUK Scotland Institute.

| Allele(s) | Model Abbreviation in Text |
| --- | --- |
| VillinCreER *BrafV600E Trp53fl/fl* | BP |
| VillinCreER *BrafV600E Rnf43fl/fl* | BRNF |
| VillinCreER *BrafV600E Trp53fl/fl Rnf43fl/fl* | BPRNF |

**Supplementary Video 1:** Time-lapse series of 3D reconstructed pure RC2 H2B-mNeon organoid in growth factor depleted medium, representative image related to Figure 2i,j.

**Supplementary Video 2:** Time-lapse series of 3D reconstructed RC2 H2B-mNeon organoid in hybrid structure with HUB040 H2B-mcherry in growth factor depleted medium, representative image related to Figure 2i,j.

**Supplementary Video 3:** Live imaging of HUB040 NDRG1-iCasp9 organoids in full medium containing 10 nM AP20187. Organoids were imaged for 16 hours and AP20187 was added right before the start of imaging. Channels are represented side by side with mNeon reporter signal on the left and brightfield on the right. Scale bar = 20 μM

